# Using modern coexistence theory to understand community disassembly

**DOI:** 10.1101/2025.08.06.668499

**Authors:** Joe Brennan, Sebastian J. Schreiber

## Abstract

Community disassembly examines how species extinction alters ecological communities. Sometimes, the extinction of one species can trigger the loss of others, known as secondary extinction. These secondary extinctions often result from complex species interactions, complicating the identification of underlying mechanisms. Here, we leverage Modern Coexistence Theory to identify when and why secondary extinctions occur. To identify when secondary extinctions occur, we introduce the community disassembly graph, that uses invasion growth rates to identify transitions between coexisting communities due to extinction. When a secondary extinction is identified, we decompose the invasion growth rates associated with the secondary extinction to understand why it occurs. We demonstrate the utility of this framework by applying it to models in which different species interactions–competition, facilitation, and predation–contribute significantly to secondary extinctions. Our results show that Modern Coexistence Theory offers a flexible and interpretable approach to understanding when and why secondary extinctions occur.

## Introduction

Species participate in a complex, entangled web of interactions, such that the extinction of one species can significantly alter the structure and dynamics of an ecological community. These cascading impacts due to species extinction, known as community disassembly, has become a central focus for understanding biodiversity change (Ostfeld and LoGiudice, 2003; Zavaleta et al., 2009). In some cases, the extinction of a single species can trigger cascading effects driving other species extinct (Paine, 1966; Lubchenco, 1978; Ebenman and Jonsson, 2005; Jordán, 2009; Langendorf et al., 2025), known as secondary extinction (Brodie et al., 2014). Secondary extinctions are speculated to contribute significantly to the loss of diversity globally under anthropogenic change in a wide variety of taxa (Moir et al., 2010; Colwell et al., 2012; Strona and Bradshaw, 2018; Kehoe et al., 2021).

When there is a strong dependency of one species on another, the cause of secondary extinction is often clear. This is the case for predators that rely on their prey or obligate mutualists that depend on their partners. Most theoretical studies of community disassembly focus on these direct dependencies. A common approach to identify secondary extinctions uses static food webs or mutualist networks where interactions between species are modeled but not changes in species densities. To study disassembly in these models, practitioners remove a node corresponding to a species and all associated edges corresponding to some species interaction and declare that species with no more resources or mutualists experience secondary extinction (Dunne et al., 2002b; Memmott et al., 2004; Bane et al., 2018). These studies have provided important insights into how the structure of ecological networks impact community disassembly (Dunne et al., 2002b, 2004; Bascompte and Stouffer, 2009; Tang et al., 2024).

Although these studies provide a level of tractability difficult to achieve in dynamic models, they neglect well-established mechanisms of secondary extinction that arise from indirect effects. For example, Paine (1966) demonstrated that the removal of the top predator *Pisaster ochraceus* from an intertidal community led to unregulated growth of the prey species *Mytilus californianus* and *Mitella polymerus*, leading to secondary extinctions of several species. These secondary extinctions would not be predicted by the static approach as *P. ochraceus* was neither a mutualist nor resource. Only by studying the strength of the interactions and associated dynamics of the community can one understand what led to the dramatic reduction in diversity after the loss of the keystone predator *P. ochraceus* (Paine, 1980; Paine et al., 2010). Secondary extinctions also occur due to the loss of other indirect factors such as ecosystem engineering (Wright et al., 2002; Yeakel et al., 2020) and intransitive competition (Jackson and Buss, 1975; Kerr et al., 2002; Edwards and Schreiber, 2010). The exclusion of these indirect factors of secondary extinction biases our understanding of community disassembly and can lead to underestimates of secondary extinctions. By incorporating direct and indirect factors into community disassembly studies, we can more accurately predict the susceptibility of ecological communities to secondary extinctions.

However, identifying the main drivers of secondary extinctions can be complicated by the cumulative impact of multiple ecological drivers and interactions between these factors (Colwell et al., 2012; Brodie et al., 2014). A secondary extinction might occur from the impact of multiple competitors or from a single dominant competitor. For example, in systems with a single limiting resource, one species alone can be sufficient to exclude others (Gause, 1934; Tilman, 1982). In contrast, microbial communities with trade-offs in growth traits can allow all pairs of species to coexist but not the full community (Manhart and Shakhnovich, 2018), implying multiple competitors are needed for exclusion of a single species. In such cases of emergent exclusion, disentangling the relative importance of ecological factors is required to understand why the secondary extinction occurred.

Modern Coexistence Theory (MCT) is an increasingly popular framework quantifying how multiple, interacting ecological factors influence coexistence outcomes. MCT’s key metric is the invasion growth rate (IGR), defined as the per capita growth rate of a species at low density while other species are at some coexistence state. MCT decomposes this IGR to identify what factors promote successful invasion of the species (Chesson, 1994; Barabás et al., 2018; Ellner et al., 2019). However, these same techniques can also explain what factors most strongly impede successful coexistence or result in invasion failure (Grainger et al., 2019a; Shoemaker et al., 2020; Spaak and Schreiber, 2023). Therefore, we can leverage MCT to quantify the factors enabling persistence of the secondarily extinct species in the full community, but lead to their exclusion in the post-extinction community.

Here, we identify when and why secondary extinctions occur using techniques from MCT. To identify when secondary extinctions occur, we use the concept of “−*i* communities” to define the community disassembly graph that describes how extinctions alter community composition. To understand why the secondary extinction occurs, we decompose and compare invasion growth rates to identify the ecological drivers responsible for the secondary extinction. We then apply this to three different models–two of which are empirically-parameterized–where secondary extinctions occur, but that static network approaches would fail to identify. Despite each case study differing in model structure and ecological factors of interest, we show that our framework can successfully classify when and why secondary extinctions occur across diverse ecological systems.

## Methods

### The community disassembly graph

To model the dynamics of ecological communities, we use deterministic differential and difference equations where *x*_*i*_(*t*) denotes the density of species *i* at time *t*. The differential equations take the form 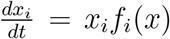 where *f*_*i*_(*x*) is the per-capita growth rate of species *I* at community state *x* = (*x*_1_, *x*_2_, …, *x*_*n*_). The discrete-time models take the form *x*_*i*_(*t* + 1) = *x*_*i*_(*t*)*F*_*i*_(*x*(*t*)) where *F*_*i*_(*x*) is the per-capita fitness at community state *x*. For discrete-time models, *f*_*i*_(*x*) = ln *F*_*i*_(*x*) corresponds to the per-capita growth rate of species *i*.

For these models, all *n* species coexist if there is a feasible attractor (Clark et al., 2024). Namely, there exists a steady state (or an invariant set) at which all species densities are positive and the species return to this steady state following sufficiently small perturbations of their densities. Similarly, for any subset of species, coexistence occurs if the model restricted to those species has a feasible attractor. One can identify these communities either analytically or numerically depending on the complexity of the model (see Appendix A).

The community disassembly graph (CDG) is a graph whose vertices correspond to subsets of coexisting species and whose edges describe transitions between these communities due to single species removals. For presentation purposes, we assume each subset of coexisting species has only one feasible attractor (see Appendix A for the general case and additional details). Let *S* ⊆ {1, 2, …, *n*} correspond to a subset of coexisting species, i.e. a vertex. Following the loss of species *i* ∈ *S* from *S*, we assume the community dynamics converge to a coexisting subset *T* ⊂ *S* without species *i*, i.e. −*i* communities of community *S* (Chesson, 1994; Hofbauer and Schreiber, 2022) (see Appendix A for dealing with convergence to a non-community attractor). Corresponding to this transition, the CDG has a directed, labeled edge 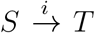. Identifying these transitions for all subsets *S* of coexisting species and the removal of any species *i* ∈ *S* from these communities defines the CDG. The CDG is a directed acyclic graph with one terminating vertex, the empty community.

### Using IGRs to identify −*i* communities and secondary extinctions

We can solve for −*i* communities using IGRs, defined as the per-capita growth rate of a species when rare and resident species are coexisting (Schreiber, 2000). For differential equations, the IGR of species *i* at some coexistence state *S* equals

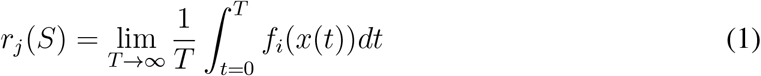

while for difference equations, they equal

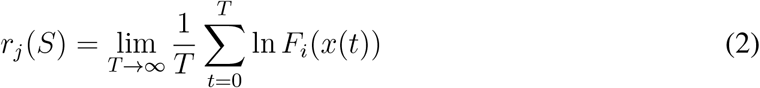

whenever the limits exist (see Appendix A).

IGRs are typically used to identify if a species can successfully invade a coexisting community. If the IGR of a species is positive, the invading species’ density will (at least initially) increase exponentially. If the IGR of a species is negative, upon introduction into the community, the species’ density will decay to extinction.

Leveraging results from Schreiber (2025) (Appendix A), one can identify −*i* communities using IGRs. Specifically, let *S* ⊆ {1, 2, …, *n*} be a subset of coexisting species. A community *T* ⊂ *S* is a −*i* community of *S* if species *i* is missing from *T* and all other missing species from *S* have negative IGRs i.e., *r*_*j*_(*T*) < 0 for *j* ∈ *S* \ {*T* ∪ {*i*}}. Intuitively, any other missing species had densities that decreased to zero. Hence, these species have negative per-capita growth rates as the system approaches community *T* i.e., *r*_*j*_(*T*) < 0. In particular, secondary extinctions occur when species in *S*, other than possibly species *i*, have a negative IGR at the −*i* community *T*.

### Using IGR decompositions to identify the cause of secondary extinction

Having identified a secondary extinction event from the community disassembly graph, we want to understand the relative importance of different ecological factors underlying this event. Therefore, we apply an MCT decomposition approach to the two IGRs associated with the secondary extinction. Say species *j* went extinct following the loss of species *i*. We know species *j* then has a negative IGR at the −*i* community. To understand why species *j* has a negative IGR at the −*i* community, we can decompose its IGR to quantify what ecological factors impede its establishment. This decomposition explains why species *j* cannot persist following the loss of species *i*.

To explain how species *i* promotes the persistence of species *j*, we identify the −*j* community of the pre-exinction community and decompose the IGR of species *j* at the −*j* community. If the pre-extinction community coexists at a global, feasible attractor, then species *j* will have a positive IGR at the −*j* community (Schreiber, 2000; Hofbauer and Schreiber, 2022; Schreiber, 2025). By decomposing the IGR of species *j* at the −*j* community, we can understand how ecological factors in the −*j* community allow species *j* to invade successfully when it could not in the −*i* community.

With both decompositions performed, we can compare them to determine what ecological factors (1) promote the persistence of species *j* in the pre-extinction community and (2) drove the exclusion species *j* following the loss of species *i*. If some factor contributes more to the IGR of species *j* in the −*j* community more than it does in the −*i* community, we can infer this factor helps species *j* persist in the presence of species *i*.

### Decomposing invasion growth rates

Given an invasion growth rate, we partition it into a sum of terms corresponding to the ecological factors of interest. Here, we use the IGR decomposition method of Ellner et al. (2019). Say species *j* is missing from a community *T*. We want to understand how *k* ecological factors (e.g., preferential predation and fluctuations of resources) affect species *j*’s IGR relative to its coexisting competitors (Shoemaker et al., 2020). Assume species *j*’s IGR for a given set of ecological factors *θ* = (*θ*_1_, …, *θ*_*k*_) is *r*_*j*_(*T, θ*) where *θ*_ℓ_ = 0 means factor ℓ is off and *θ*_ℓ_ = 1 means it is on. If we are interested in first-order and interactive effects of these factors on the invasion growth rate, we decompose the invasion growth rate at a feasible attractor supporting only the species in *T* as follows:

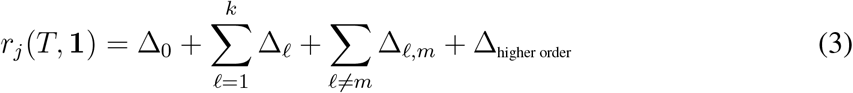

where **1** = (1, 1, …, 1). Δ_0_ is the baseline value relative to the average influence on resident competitor species’ IGRs when all factors are off. Specifically,

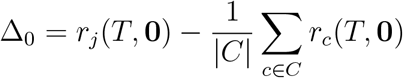

where **0** = (0, 0, …, 0) and *C* ⊆ *T* is the set of resident competitor species. Biologically, Δ_0_ represents the difference in the average per-capita growth rate of species *j* and the resident competitors when all of the ecological factors of interest are turned off.

Δ_ℓ_ is the change in the IGR relative to the average influence on resident competitor species’ IGRs when only factor ℓ is on. Specifically,

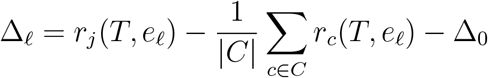

where *e*_ℓ_ is the vector with 1 in entry ℓ and otherwise zero. When Δ_ℓ_ > 0, the ecological factor is more beneficial (or less harmful) to the focal species than its average impact on competitors. Conversely, if Δ_ℓ_ < 0, the ecological factor is less beneficial (or more harmful) to the focal species than its average impact on competitors.

Regarding interactive effects between ecological factors, Δ_ℓ,*m*_ is the change in the IGR relative to the average resident competitors’ IGRs due to the interactive effects of factors ℓ ≠ m. Specifically,

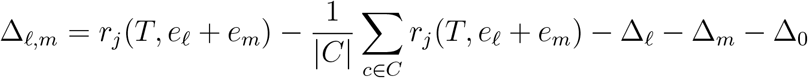

There will be a non-zero interaction term if the two factors have a nonlinear relationship in the invasion growth rates.

Lastly, Δ_higher order_ are higher order effects. Specifically,

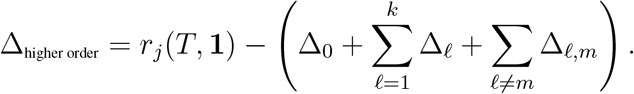

## Applications

We employ our framework in three different models, each in which different species interactions help maintain coexistence of the full system: competition, facilitation, and predation. In each of these cases, the species interaction contributes significantly to the persistence of the full community, and thus, following loss of the species mediating this interaction, another species experiences extinction. Crucially, each of these examples highlights a secondary extinction that cannot be characterized by classic, static approaches to secondary extinctions.

### Competition

Historically, competition has been viewed as a species interaction that disrupts coexistence (Armstrong and McGehee, 1980; Pastore et al., 2021). However, competition can also promote coexistence by suppressing the densities of competitively dominant species, allowing otherwise inferior competitors to persist (Edwards and Schreiber, 2010; Levine et al., 2017). To illustrate an example of competition-mediated coexistence, we analyze an empirically parameterized model of a competitive annual plant community (Van Dyke et al., 2022).

Van Dyke et al. (2022)’s model tracks annual seed production. The seeds germinate with probability *g*_*i*_ or go into the seed bank where they survive with probability *s*_*i*_ to the next year. The germinated seeds compete and then produce seeds. Under these assumptions, Van Dyke et al. (2022)’s model is

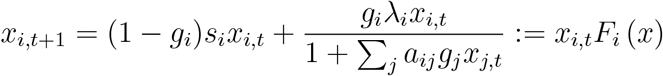

For illustrative purposes, we focus on the model parametrized to the ambient rain scenario for the largest subset of coexisting species: *Acmispon wrangelianus, Hordeum murinum*, and *Plantago erecta* (parameters in Appendix B). To create the community disassembly graph for this model, we find all coexistence states analytically (see Appendix B). These correspond to all single species communities, a single species pair (*A. wrangelianus* and *H. murinum*), and all three species.

Next, we evaluate each species’ IGR at the single species and two species coexistence states. Each of these corresponds to a stable equilibrium 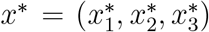. The IGR of species *i* at these equilibria equal

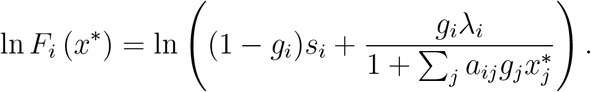

Using these IGRs, we construct the community disassembly graph (Figure 1A).

**Figure 1:**
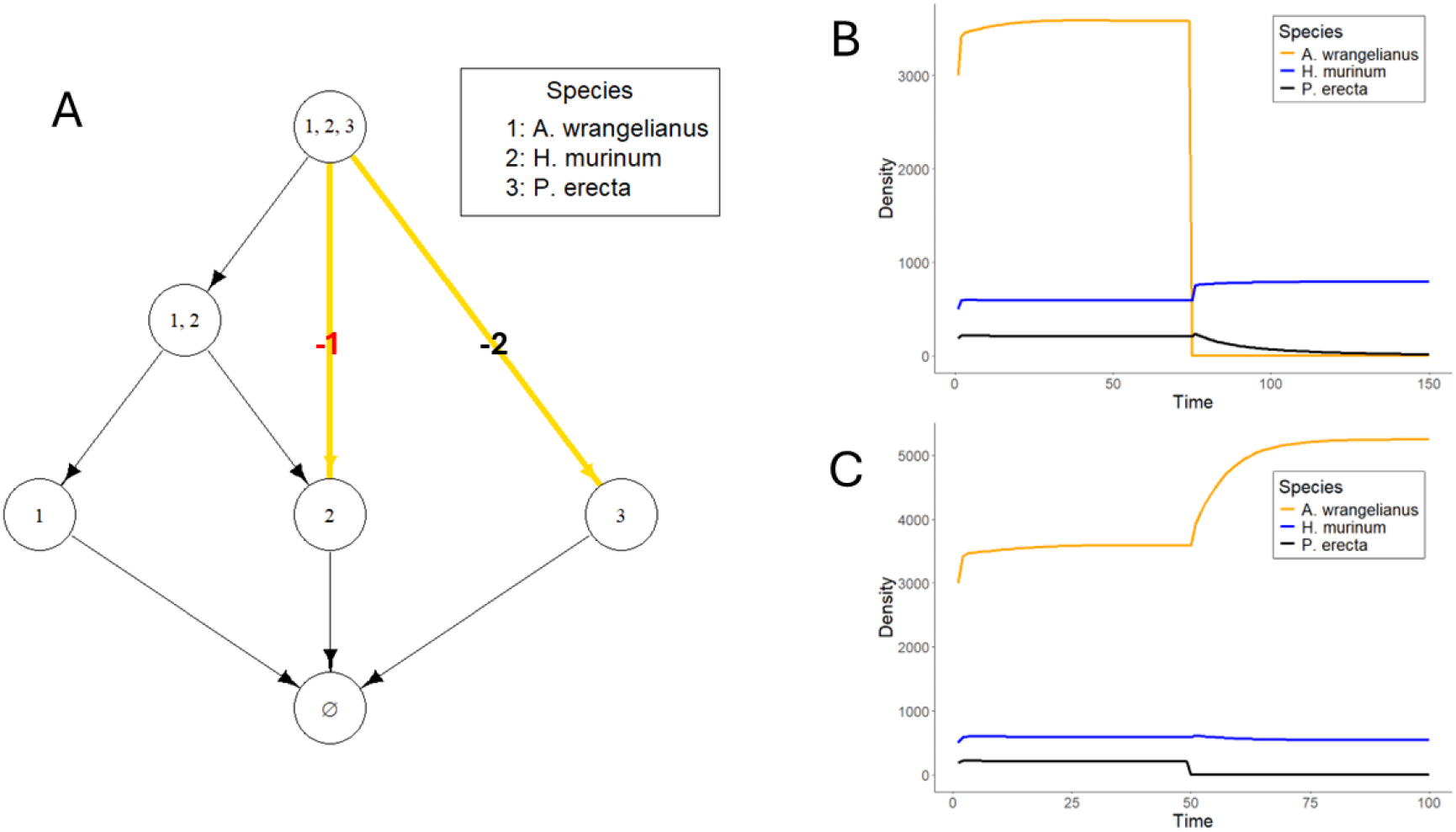
(A) Community disassembly graph of the annual plant community. Vertices depict coexisting communities. Directed edges represent transitions from one community to another due to extinction of the labeled species. Extinctions resulting in no secondary extinctions are denoted by black directed edges with no edge label while those containing secondary extinctions are in yellow with a corresponding edge label. The analyzed secondary extinction event is denoted by a red edge label. (B) Simulation of community dynamics following extinction of *P. erecta* at time step 50. Note that there are no secondary extinction events, matching the -*P. erecta* community predicted in the community disassembly graph. (C) Simulation of community dynamics following extinction of *A. wrangelianus* at time step 75. *P. erecta* decays towards extinction following the extinction of *A. wrangelianus*, confirming the secondary extinction predicted on the community disassembly graph.

This community disassembly graph identifies two secondary extinction events: the secondary extinction of *A. wrangelianus* following the loss of *H. murinum* and the secondary extinction of *P. erecta* following the loss of *A. wrangelianus* (Figure 1A). We focus here on understanding why *P. erecta* goes extinct following the extinction of *A. wrangelianus* (Figure 1B). Appendix C analyzes the other case. To determine the cause of the secondary extinction, we examine (1) what factors prevent *P. erecta* from establishing when only *H. murinum* is present and (2) what factors allow *P. erecta* to invade when *A. wrangelianus* is present in the community with *H. murinum*. Specifically, the factors of interest are competition against *P. erecta* and indirect impacts of competition between *A. wrangelianus* and *H. murinum* on *P. erecta*. Following equation 3, we have *k* = 2 where *θ*_1_ = 0 denotes setting all interspecific competition coefficients against *P. erecta* to zero while *θ*_2_ = 0 denotes setting all interspecific competition coefficients against *H. murinum* and *A. wrangelianus* to zero. Because these factors are additive in the invasion growth rates, the higher order terms Δ_ℓ,*m*_ are zero.

*P. erecta* has a negative IGR at the *H. murinum* community (i.e., the −*A. wrangelianus* community), as it undergoes a secondary extinction following removal of *A. wrangelianus* (Figure 1A & B). Our decomposition of this IGR (Figure 2) shows, as expected, the direct impact of competition from *H. murinum* on *P. erecta* is why this IGR is negative.

**Figure 2:**
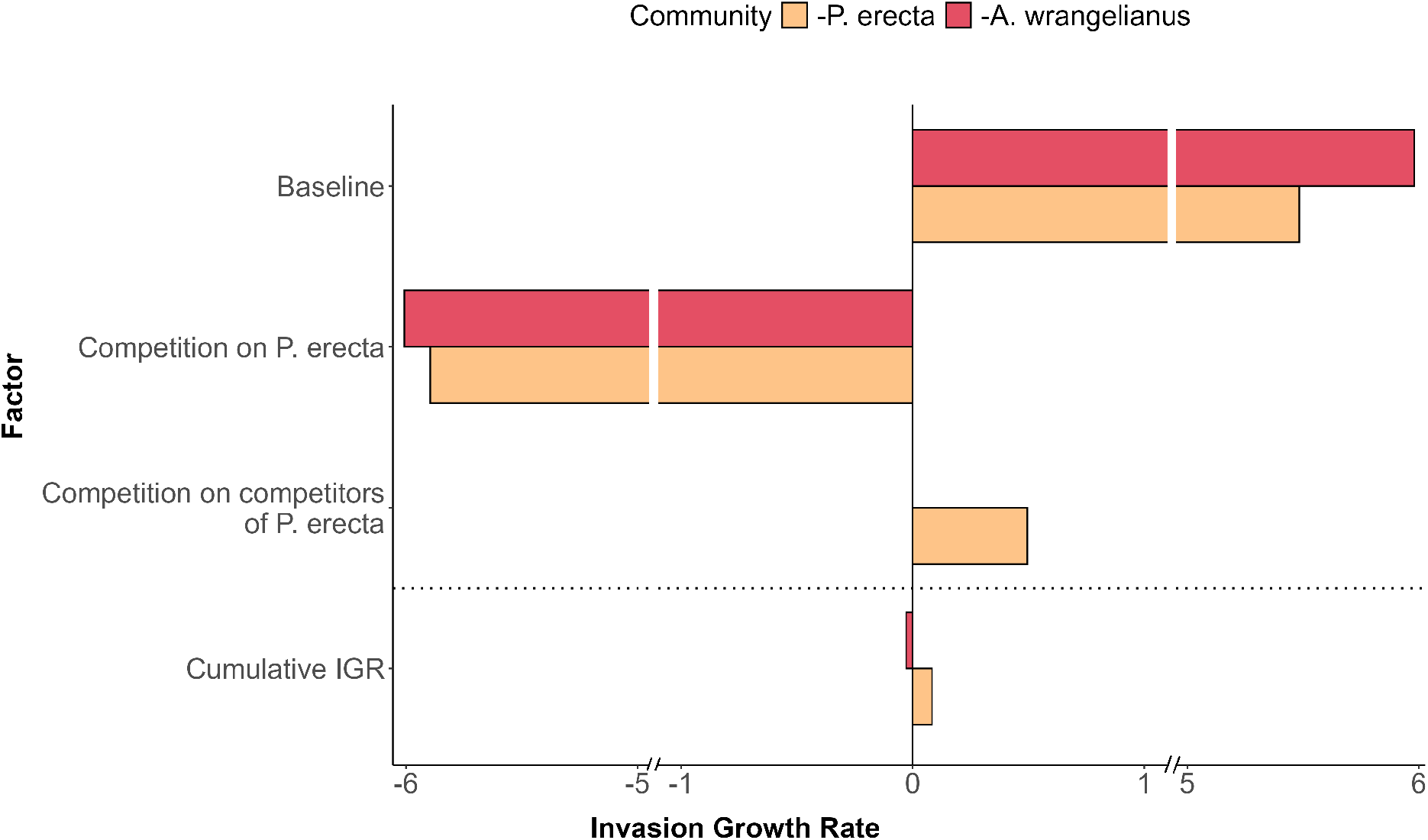
Decomposition of *P. erecta*’s IGR at the −*P. erecta* and −*A. wrangelianus* communities. The sum of all bars of a respective color above the dotted line add to the value of the bar below the dotted line of the respective color. Factors not present in the −*i* community have no contribution to the IGR and thus have no corresponding bar.

To understand why *P. erecta* can invade with *H. murinum* only in the presence of *A. wrangelianus*, we decompose its IGR at the *A. wrangelianus*–*H. murinum* community i.e., the −*P. erecata* community (Figure 1A & C). In this community, the baseline conditions become less beneficial to *P. erecta*. Conversely, we find competition on *P. erecta* becomes weaker in the presence of *A. wrangelianus*. Importantly, competition between *A. wrangelanius* and *H. murinum* has a positive impact on the IGR of *P. erecta*. The positive contribution of competition between *P. erecta*’s two competitors drives the IGR to be positive, allowing for successful invasion and coexistence.

This analysis identifies the cause of *P. erecta*’s secondary extinction. In the community with *H. murinum* and *A. wrangelianus*, competition between *H. murinum* and *A. wrangelianus* suppresses their competitive dominance over *P. erecta*, permitting *P. erecta* to coexist. Consequently, the extinction of *A. wrangelianus* removes this competitive suppression of *H. murinum*, allowing *H. murinum* to competitively exclude *P. erecta*.

### Facilitation

Facilitation can buffer species’ densities against negative interactions such as predation (Glynn, 1976; Stachowicz and Hay, 1999) and competition (Pennings and Callaway, 1996; Stachowicz, 2001). However, dependence on facilitative or mutualist interactions can increase vulnerability to secondary extinctions (Aslan et al., 2013; Vanbergen et al., 2017).

For example, Geijzendorffer et al. (2011) empirically parametrized a series of Lotka-Volterra models to a grassland system exhibiting competitive and facilitative interactions. In these models, each species *i* has an intrinsic rate of growth, *r*_*i*_, and a per-capita impact *a*_*ji*_ on species *j*. If *a*_*ji*_ is negative, species *i* has a competitive effect on species *j* while if positive, species *I* facilitates species *j*. The model is

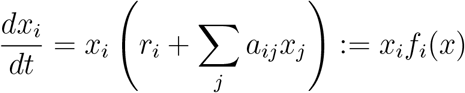

We focus on the parameterization for spring in Appendix E and consider one of two largest subsets of five coexisting species: *Agrostis stolonifera, Lolium perenne, Phleum pratense, Trifolium pratense*, and *Trifolium repens*.

Using the invasion graph method (Hofbauer and Schreiber, 2022), we identify all coexistence states (Appendix E). Next, we evaluate the IGR of each species at each coexistence state. Due to the time averaging property of Lotka-Volterra models (Hofbauer and Sigmund, 1998), these IGRs correspond to evaluating the per-capita growth rates at the respective equilibrium 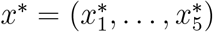

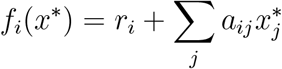

Using this information, we construct the community disassembly graph (Figure 3A). This graph reveals eight secondary extinction events. Half of the secondary extinction events occur following the extinction of *T. repens*, causing the secondary extinction of *T. pratense*. This suggests *T. repens* plays a crucial role in allowing *T. pratense* to coexist with other species in multiple contexts.

**Figure 3:**
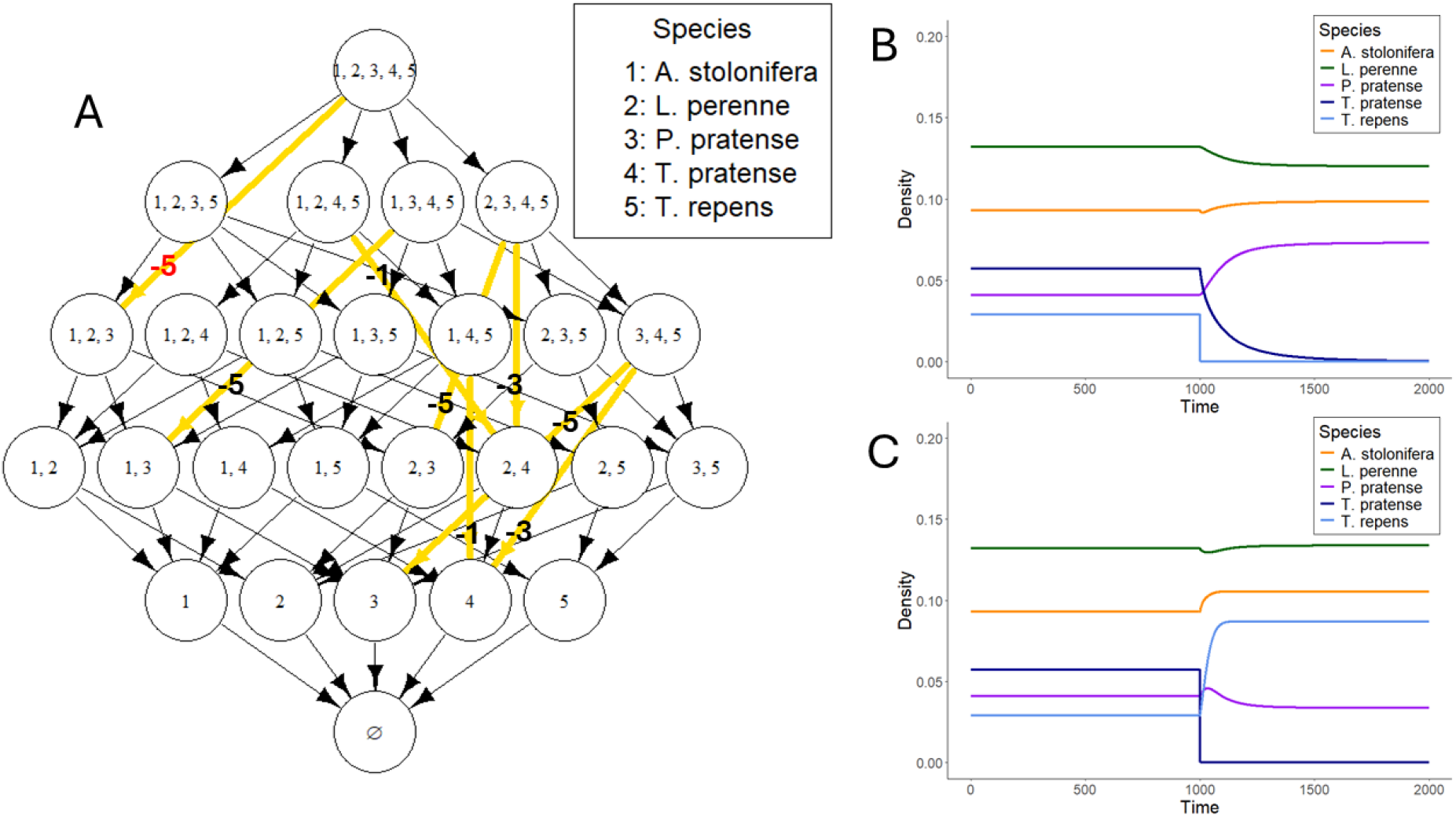
(A) Community disassembly graph of the grassland community. Vertices depict coexisting communities. Directed edges represent transitions from one community to another due to extinction of the labeled species. Extinctions resulting in no secondary extinctions are denoted by black directed edges with no edge label while those containing secondary extinctions are in yellow with a corresponding edge label. The secondary extinction event whose mechanisms are quantified is denoted by a red edge label. (B) Simulation of community dynamics following extinction of *T. repens* at time step 1, 000. *T. pratense* decays towards extinction following the extinction of *T. repens*, confirming the secondary extinction predicted on the community disassembly graph. (C) Simulation of community dynamics following extinction of *T. pratense* at time step 1, 000. Note that there are no secondary extinction events, matching the -*T. pratense* community predicted in the community disassembly graph.

To understand these secondary extinctions, we focus on the secondary extinction of *T. pratense* when *T. repens* is removed from the entire community (Figure 3A & B). To understand why this secondary extinction event occurs, we address two key questions: (1) what factors prevent *T. pratense* from invading the community with *A. stolonifera, L. perenne*, and *P. pratense* (i.e., the -*T. repens* community), and (2) what factors allow *T. pratense* to invade the community with these species if *T. repens* is present (i.e., the −*T. pratense* community)? Specifically, we quantify the contributions of the direct effects of competition and facilitation on *T. pratense* and the indirect effects of competition and facilitation on competitors of *T. pratense*. Following equation 3, we have *k* = 4 where *θ*_1_ = 0 and *θ*_2_ = 0 denote setting all competitive and facilitative interactions against *T. pratense* respectively to zero, while *θ*_3_ = 0 and *θ*_4_ = 0 denote setting all competition and facilitative interactions against competitors of *T. pratense* respectively to zero. Because these factors are additive in the invasion growth rates, the higher order terms Δ_ℓ,*m*_ are zero.

To understand why *T. pratense* cannot establish itself in the −*T. repens* community, we decompose its negative IGR when the other species are at equilibrium. We find that direct effects of competition and indirect effects of facilitation among competitors outweigh the indirect effects of competition among competitors. This results in a negative IGR for *T. pratense*. However, without the indirect effect of facilitation among competitors, the IGR would be positive. Hence, facilitation among competitors plays a crucial role in the competitive exclusion of *T. pratense*.

Next, we examine how the presence of *T. repens* allows the establishment of *T. pratense* i.e., in the −*T. pratense* community (Figure 3A & C). Unlike in the −*T. repens* community, direct facilitation on *T. pratense* confers a positive contribution to its IGR. Additionally, competition against *T. pratense* is less harsh in the −*T. pratense* community. In the −*T. pratense* community, indirect effects due to competition and facilitation are stronger. However, since they have opposing effects on the IGRs of the same magnitude, the net impact of these indirect effects are negligible on *T. pratense*’s IGR. These factors allow for *T. pratense* to have a positive IGR in the −*T. pratense* community unlike in the −*T. repens* community.

These decompositions imply that the secondary extinction of *T. pratense* following the loss of *T. repens* was primarily driven by the loss of facilitation and stronger competitive pressure. In the full community, *T. repens*’s facilitation permits *T. pratense* to persist despite competitive pressures. However, following the extinction of *T. repens, T. pratense* becomes unable to withstand competition from the remaining species in the disassembled community due to the stronger direct competition against *T. pratense* and the indirect effect of facilitation among its competitors.

### Predation

Predation can maintain coexistence by regulating competitively dominant prey species, known as keystone predation (Paine, 1966; Letnic et al., 2009; Wallach et al., 2015). A well-studied model of keystone predation is the diamond model with a basal resource (*R*), two competitors (*C*_1_ and *C*_2_) competing for the basal resource, and a top predator (*P*) that predates competitors (McCann et al., 1998; Schreiber and Rittenhouse, 2004; Vasseur and Fox, 2007; Shoemaker et al., 2020). Following Vasseur and Fox (2007)’s model formulation and parameterization in Appendix F, the dynamics are

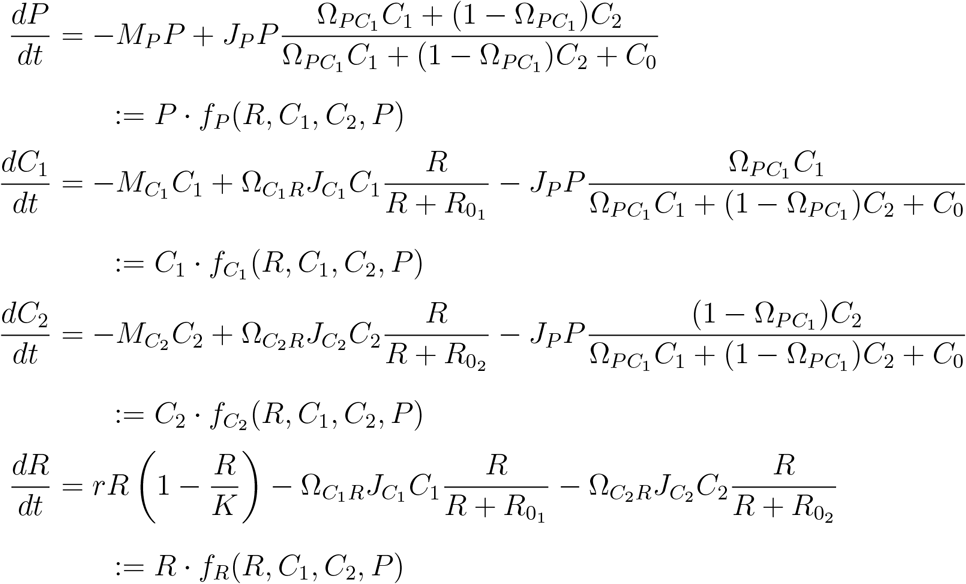

We used numerical simulations to identify coexistence states involving two or more species. For each subset containing two or more species, we ran 250 simulations with random initial densities for each species drawn uniformly between 0 and 2. Simulations were run until the dynamics reached steady state. Coexistence states were identified as sets of species with mean densities greater than 10^−9^ from the end of the burn-in period to the end of the time series.

Across all coexisting subsystems, we identified at most one attractor.

To calculate the IGR for a given coexistence state, we numerically integrated the model using the deSolve package in the programming language R (Soetaert et al., 2010) evaluated at time intervals of Δ*t* = 0.1 with a burn-in time of 1, 000 and a total run time of 10, 000. We approximate the IGR for species *i*

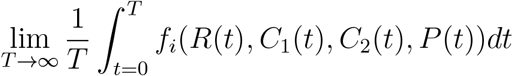

by the Riemann sum

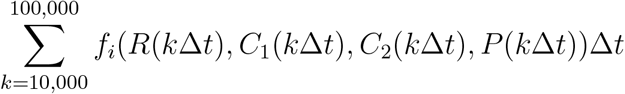

for *i* ∈ {*R, C*_1_, *C*_2_, *P* }.

Using the numerically identified coexistence states and associated IGRs, we construct the community disassembly graph (Figure 5). We find several secondary extinctions (Figure 5A). Most of the secondary extinctions are intuitive (loss of resource leads to loss of species directly dependent on the resource). However, two are more interesting: extinction of the predator *P* induces secondary extinction of the second consumer *C*_2_ and the extinction of the first consumer *C*_1_ induces the secondary extinction of the predator *P*. Here, we study how the extinction of *P* induces the secondary extinction of *C*_2_ (Figure 5A & B). We consider three ecological factors: fluctuations in resource density, predator preference 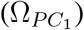, and the resident consumer’s utilization of the shared resource 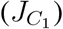. Following equation 3, we have *k* = 3 where *θ*_1_ = 0 denotes setting resource density to its average, *θ*_2_ = 0 denotes eliminating predator preference 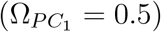, and *θ*_3_ = 0 denotes eliminating the resident consumer’s utilization of the shared resource 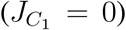. All of these factors are additive in the invasion growth rates except for fluctuations in resource density and the resident consumer’s utilization of the shared resource. Thus, we will have Δ_0_, Δ_ℓ_ for each factor, and only have the interaction terms Δ_ℓ,*m*_ for fluctuations in resource density and the resident consumer’s utilization of the shared resource.

**Figure 4:**
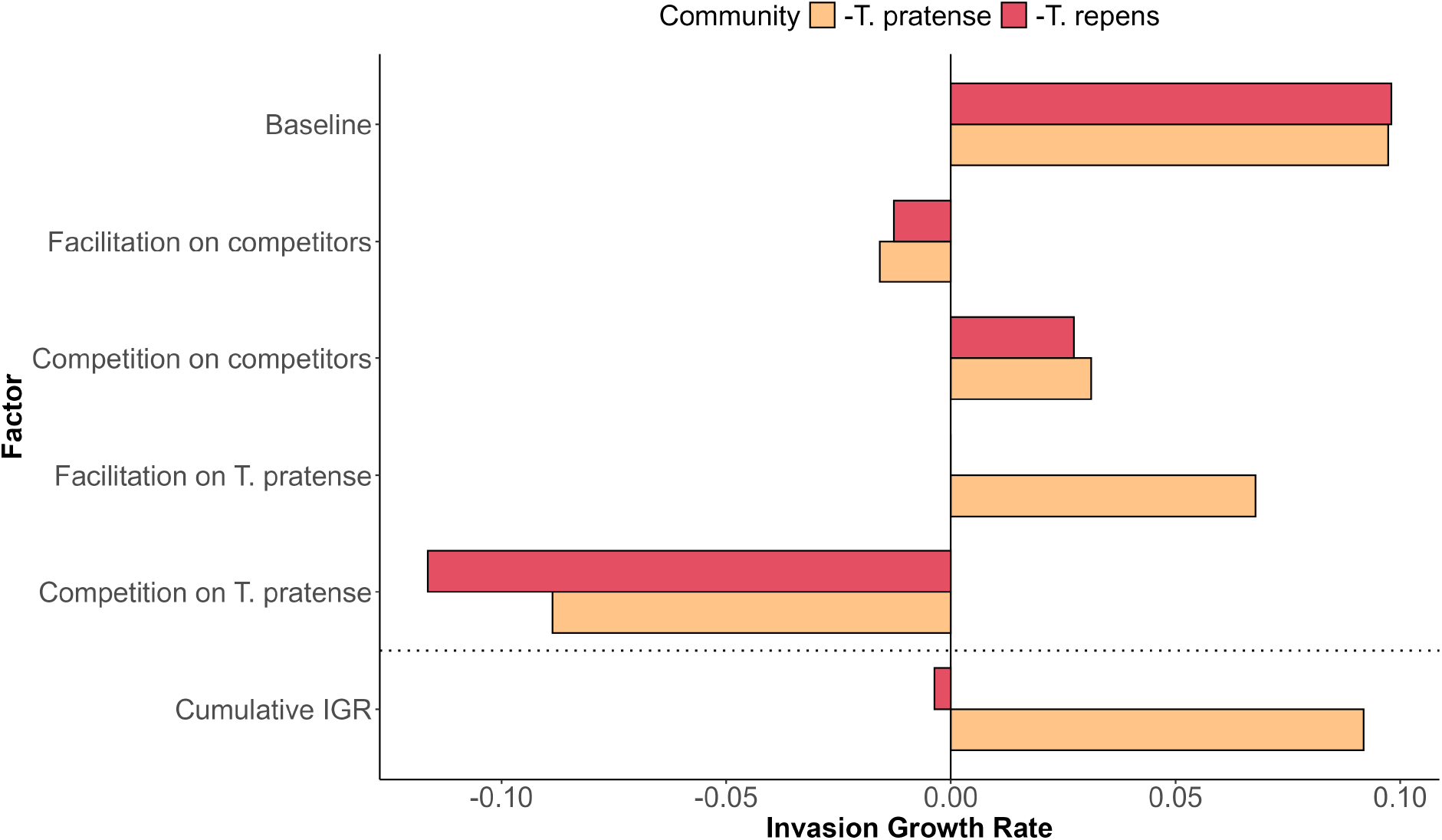
Decomposition of *T. pratense*’s IGR at the −*T. pratense* and −*T. repens* communities. The sum of all bars of a respective color above the dotted line sum to the value of the bar below the dotted line of the respective color. Factors not present in the −*i* community have no contribution to the IGR and thus have no bar.

**Figure 5:**
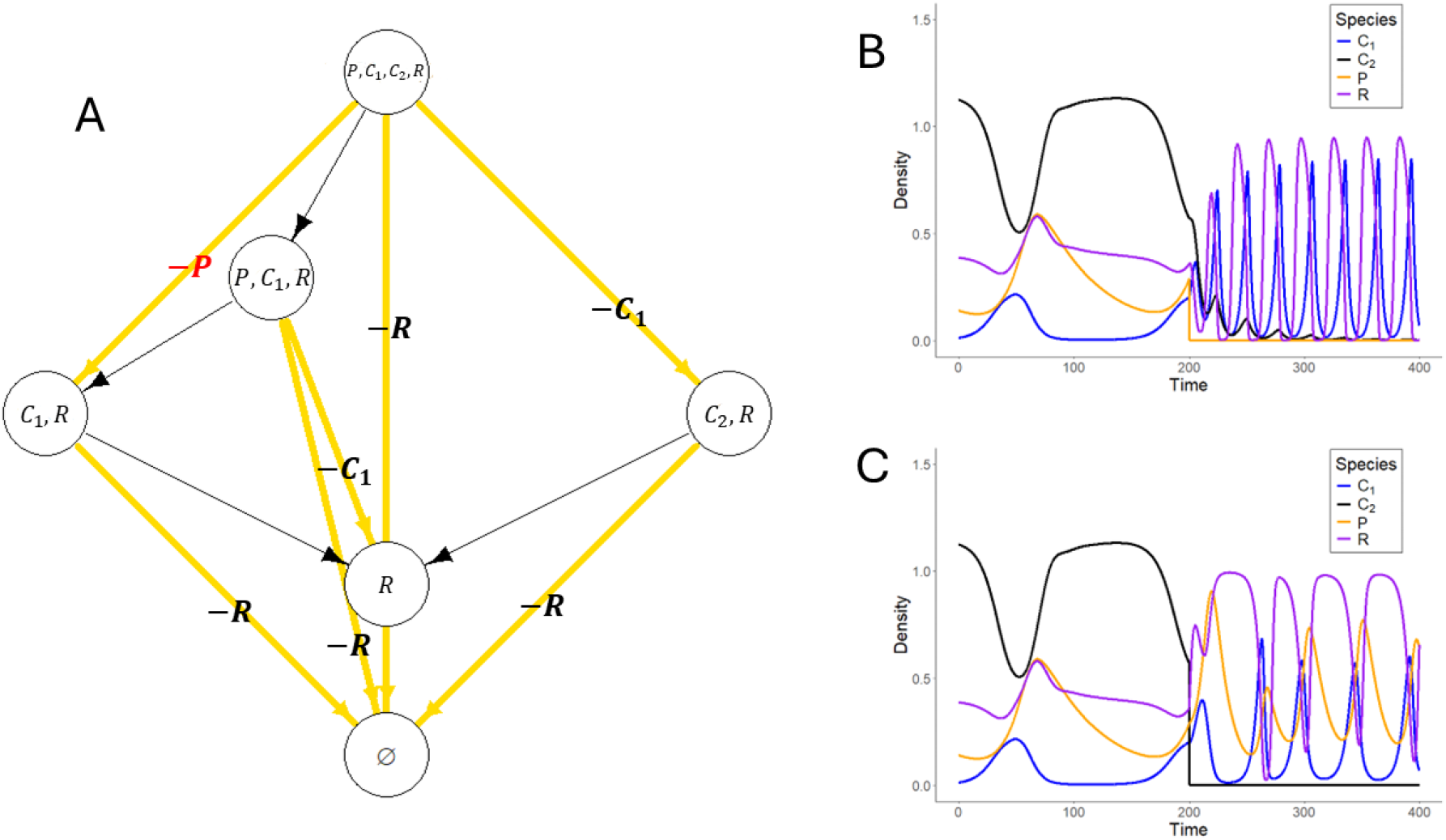
(A) Community disassembly graph of the diamond model. Vertices depict coexisting communities. Directed edges represent transitions from one community to another due to extinction of the labeled species. Extinctions resulting in no secondary extinctions are denoted by black directed edges with no edge label while those containing secondary extinctions are in yellow with a corresponding edge label. The secondary extinction event whose mechanisms are quantified is denoted by a red edge label. (B) Simulation of community dynamics following extinction of *P* at time step 200. *C*_2_ decays towards extinction following the extinction of *P*, confirming the secondary extinction predicted on the community disassembly graph. (C) Simulation of community dynamics following extinction of *C*_2_ at time step 200. Note that there are no secondary extinction events, matching the −*C*_2_ community predicted in the community disassembly graph.

First, we assess why consumer *C*_2_ has a negative IGR in the community with only the resource *R* and the other consumer *C*_1_ i.e., the −*P* community (Figure 5A & B). As expected, the utilization of the resource by *C*_1_ is the primary factor that makes *C*_2_’s IGR negative (Figure 6). We find fluctuations in *R* have a negative impact on the IGR of *C*_2_, but the interaction between fluctuations in *R* and *C*_1_’s intake of *R* has a positive impact on *C*_2_’s IGR due to its negative impact on *C*_1_. This positive contribution, however, does not offset the negative contributions of each factor independently, resulting in a negative IGR of *C*_2_.

**Figure 6:**
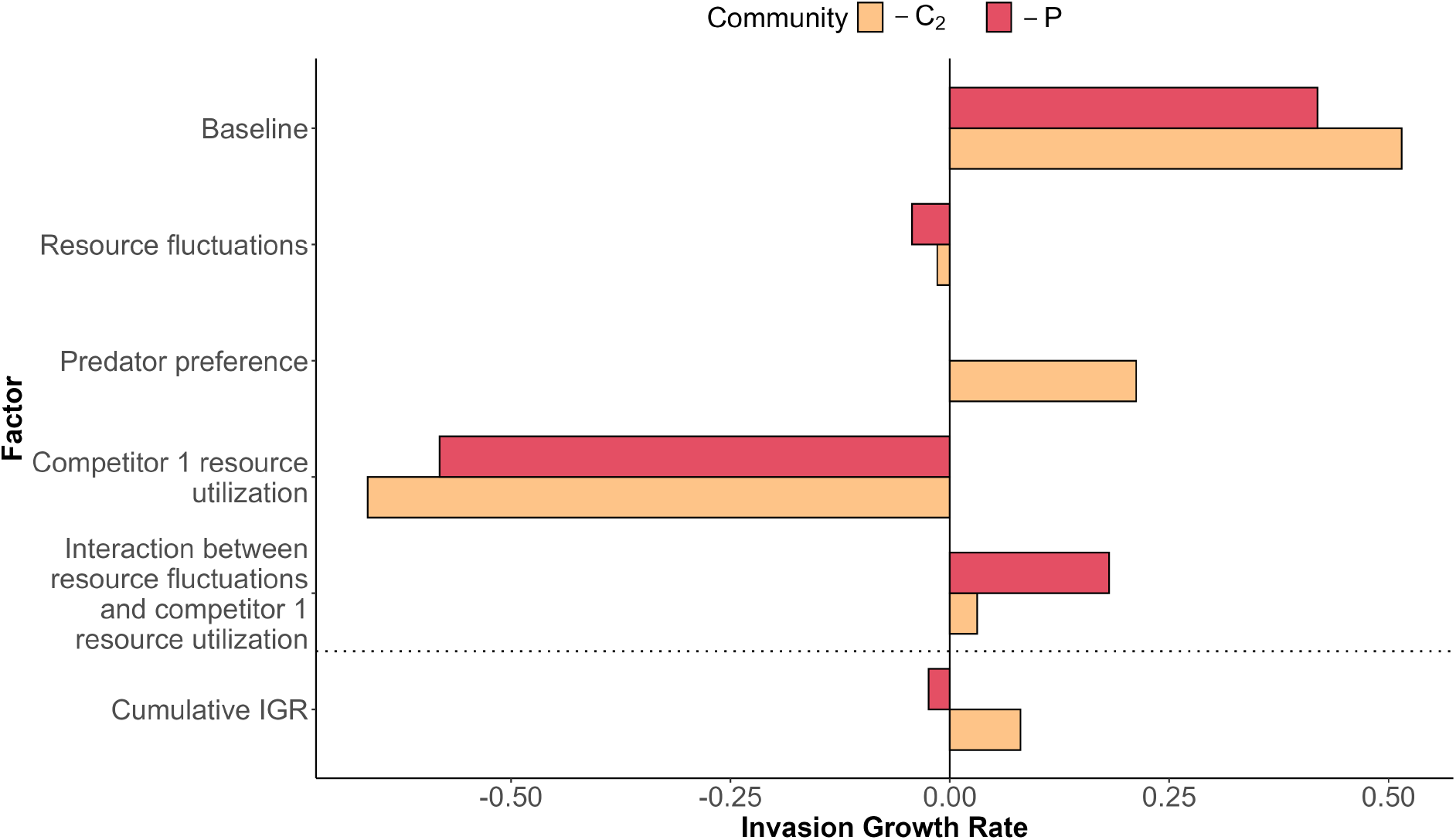
Decomposition of *C*_2_’s IGR at the −*C*_2_ and −*P* communities. The sum of all bars of a respective color above the dotted line add to the value of the bar below the dotted line of the respective color. Factors not present in the −*i* community have no contribution to the IGR and thus have no bar represented.

Next, we examine why *C*_2_ can establish in the presence of the predator *P*. To do this, we decompose the IGR of *C*_2_ at the −*C*_2_ community (Figure 5A & C). Decomposing the IGR of *C*_2_, we find fluctuations in *R* and *C*_1_’s use of *R* have negative contributions to the IGR while their interaction has a weak, positive contribution (Figure 6). The effect of predator preference is sufficiently positive to counteract the negative contributions of fluctuations in *R* and *C*_1_’s utilization of *R*.

Comparing the two IGR decompositions, we find the predator’s presence made baseline conditions and resource fluctuations more beneficial for *C*_2_ while making the impact of *C*_1_’s resource utilization and the interaction between these factors more detrimental for *C*_2_. In addition to changing these pre-existing factors, the predator’s preference serves as a positive factor resulting in a positive IGR.

Synthesizing these results, when *P* is present, its preferential consumption of *C*_1_ suppresses *C*_1_’s competitive dominance over *C*_2_, enabling coexistence of all species. However, the extinction of *P* eliminates the top down control on *C*_1_. Without this top-down control, *C*_1_’s utilization of *R* made it a superior competitor, driving competitive exclusion of *C*_2_. Thus, *C*_2_’s coexistence in the full community depends on *P*’s presence, implying extinction of *P* would drive a subsequent loss of *C*_2_.

## Discussion

While invasion growth rates (IGRs) have traditionally been used to identify mechanisms of species coexistence, they are increasingly being applied to more ecological phenomena, including priority effects (Ke and Letten, 2018; Grainger et al., 2019a), competitive exclusion (Germain et al., 2018; Shoemaker et al., 2020), and community assembly (Spaak and Schreiber, 2023; Song and Spaak, 2024). We contribute to this expanding body of work by extending key concepts from Modern Coexistence Theory such as IGRs and −*i* communities to the study of community disassembly. In doing so, we use IGRs to identify when and why secondary extinctions occur, further demonstrating their utility beyond coexistence analysis (Grainger et al., 2019b).

Our approach to community disassembly relies on an underappreciated cornerstone of Modern Coexistence Theory: −*i* communities. These communities, introduced by Chesson (1994), correspond to the coexisting species remaining in the long-term after species *i* is lost. The community disassembly graph introduced here uses this concept recursively, thus describing the cascading effects of species removals. As illustrated in our applications, secondary extinctions can lead to −*i* communities where multiple species, not only species *i*, are missing. However, almost all theoretical developments and applications of IGRs focus on species whose loss does not lead to secondary extinctions (MacArthur and Levins, 1967; Chesson, 1994; Ellner et al., 2019; Shoemaker et al., 2020; Spaak and Schreiber, 2023). This limitation suggests MCT frameworks must be expanded to account for systems where species losses trigger cascading extinctions.

Community disassembly graphs reveal mechanisms of secondary extinction complementing insights from static network analyses, structural coexistence frameworks, and large random community models. Static approaches capture bottom-up extinctions after resource loss and the loss of obligate mutualists (Dunne et al., 2002b; Vanbergen et al., 2017), but miss secondary extinctions driven by indirect competition, non-obligate facilitation, and keystone predation. Dynamical food-web analyses capture bottom-up and top-down cascades and show how disruption of predator-mediated coexistence can trigger secondary extinctions (Eklöf and Ebenman, 2006; Curtsdotter et al., 2011). Our approach extends this by demonstrating how competitive and facilitative interactions drive secondary extinctions including indirect facilitation where species loss releases stronger competitors (Figure 2) and a combination of direct facilitation and indirect effects (Figure 4). This complements random matrix approaches predicting the statistical properties of post-extinction abundances in large model ecosystems (Rossberg et al., 2017; Emary and Evans, 2021). Similarly, structural approaches to coexistence in empirical communities demonstrate multi-trophic interactions can allow coexistence otherwise not possible within a single trophic level (Saavedra et al., 2017; Bartomeus et al., 2021). This implies the extinction of higher trophic level species could drive secondary extinctions, analogous to our findings in the diamond model (Figure 6). These approaches are thus complementary: random matrix theory provides statistical expectations for large systems, structural approaches meld empirical parameterization and theoretical parameter exploration, and disassembly graphs reveal the details that manifest those statistical patterns in real communities.

The applications of these methods demonstrate secondary extinctions can arise from multiple factors. For example, in the grassland system (Figure 4), the secondary extinction of *T. pratense* was driven by increased direct competition and the loss of direct facilitation in the disassembled community. Similarly, in the annual plant model, an alternate decomposition demonstrates both the interactive effect of competition from *A. wrangelianus* and *H. murinum* on *P. erecta* and competition between *H. murinum* and *A. wrangelianus* are needed to allow successful invasion and coexistence of *P. erecta* (Appendix D). More broadly, these results highlight how secondary extinctions can be driven by complex combinations of direct and indirect interactions that are disrupted when species are lost from communities, supporting previous studies suggesting ecological factors and stressors can interact in additive and synergistic ways (Graham et al., 2011; Colwell et al., 2012; Brodie et al., 2014).

## Challenges and future directions

Beyond identifying when secondary extinctions occur, the community disassembly graph provides a framework for addressing a broader set of questions about how communities change structure during species loss. Because it tracks all possible changes in community composition arising from extinction events, it enables the analysis of how multi-species extinction sequences influence community dynamics. One example is how the order and timing of extinctions shape the composition of disassembled communities, known as inverse priority effects (Torres et al., 2024a,b). Additionally, future studies could investigate how the topology and strength of species interactions influence the structural properties of the community disassembly graph (Bascompte and Stouffer, 2009; Zavaleta et al., 2009).

Although these applications highlight the potential of community disassembly graphs, their implementation faces important computational and data limitations. They require substantially more data than static approaches because of their reliance on explicitly modeling community dynamics. Moreover, as ecological complexity increases, computing community disassembly graphs becomes increasingly computationally intensive: there are 2^*n*^ possible coexistence states for a pool of *n* species, making exhaustive enumeration infeasible for more speciose communities. However, in many applications, interest centers on secondary extinctions for a particular community of *n* coexisting species (Eichenwald and Reed, 2021). In such cases, rather than identifying all feasible attractors across all richness levels, one only needs to identify attractors for the *n* subsystems consisting of *n* − 1 species i.e., all of the −*i* communities of *n* species community. This targeted approach substantially reduces computational demands while still addressing the key question of community robustness to single-species losses.

Our framework assumes following a species’ removal, community dynamics converge to a feasible attractor of the remaining species. However, the removal of a species can instead cause heteroclinic cycling, where the trajectory approaches a sequence of different feasible attractors, spending progressively longer times near each before transitioning to the next (May and Leonard, 1975; Hofbauer, 1994; Schreiber and Rittenhouse, 2004; Song and Spaak, 2024). When such intransitive attractors occur, the disassembly graph must be expanded to account for reaching any community along the cycle (see Appendix A). However, identifying these intransitive attractors using IGRs presents a substantial mathematical challenge (Hofbauer, 1994). Similarly, decomposing invasion growth rates along these cycles to understand the mechanisms driving community transitions exist only for the three-species rock-paper-scissors dynamics (Ranjan et al., 2024). Extending these methods to longer cycles or more complex interaction structures remains an open challenge.

As in other studies of community disassembly, we assume instantaneous species extinction when identifying disassembled communities (Dunne et al., 2002a; Ebenman et al., 2004; Ebenman and Jonsson, 2005). Although this assumption is common, ecological dynamics may differ substantially when species instead decline gradually, as this decline can trigger indirect effects causing other species to go extinct first or shift the community into a different basin of attraction when alternative stable states exist (Hastings et al., 2018; Abbott et al., 2024). The community disassembly graph applies most directly when extinction occurs rapidly relative to other ecological dynamics or when the factors causing extinction directly affect only the focal species. In cases where gradual environmental change strongly influences multiple species, the community disassembly graph becomes less informative, since the key challenge is understanding how the feasible attractors change with the environmental conditions. However, after an extinction occurs beyond a critical environmental threshold, performing an IGR decomposition before and after the environmental threshold can identify the relative importance of shifting environmental conditions and other ecological factors.

MCT relies on species having positive invasion growth rates when rare in resident communities to ensure coexistence. However, several ecological phenomena prevent this condition from holding. For example, species exhibiting Allee effects have negative growth rates at low densities (Barabás et al., 2018), while obligate mutualists have negative invasion growth rates in communities lacking their partners. In our framework, the presence of Allee effects or obligate mutualisms does not prevent the identification of secondary extinctions. However, our IGR decomposition becomes uninformative in explaining why such species go extinct, because the species cannot reinvade through low-density introductions and therefore cannot return to the pre-extinction community (Spaak and Schreiber, 2023). Although frameworks exist to characterize coexistence in some of these cases (Schreiber et al., 2019; Ellner et al., 2022), the analog of an IGR decomposition to identify the causes of secondary extinctions in communities with Allee effects or obligate mutualisms remains an open challenge.

By applying MCT to community disassembly, we provide both a framework for identifying secondary extinctions–the community disassembly graph–and a method for understanding their underlying causes through IGR decomposition. Our approach reveals that understanding why species go secondarily extinct requires analyzing invasion growth rates at multiple communities to determine both how species coexist initially and why they fail following primary extinctions. This work demonstrates invasion growth rates provide a powerful lens not only for understanding species coexistence, but also for predicting and explaining community responses to species loss.

## Acknowledgments

We thank Marissa Baskett for comments on an early draft. We also thank Raissa D’Souza, Alan Hastings, Marcel Holyoak, and John Stachowicz for feedback throughout the development of this project. We thank Mary Van Dyke and Nathan Kraft for discussions about the annual plant system. We also thank four anonymous reviewers for thoughtful reviews that enhanced the quality of the manuscript. JB was supported by the National Science Foundation Graduate Research Fellowship program under Grant No. DGE-2439024.

## A The community disassembly graph: Mathematical and Numerical Considerations

In this Appendix, we present the general definition of the community disassembly graph, the mathematical assumptions underlying this definition, and the characterization of directed edges of this graph using invasion growth rates. We also discuss the implications of our underlying assumptions for numerically identifying coexisting sets of species (the vertices) and the associated invasion growth rates used to identify the edges.

Our presentation focuses on continuous-time models 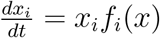 where *x* = (*x*_1_, *x*_2_, …, *x*_*n*_) is the vector of species densities, and *f*_*i*_ is the per-capita growth rate of species *i*. However, all the definitions, assumptions, results and numerical considerations carry over to the difference equations *x*_*i*_(*t* + 1) = *x*_*i*_(*t*)*F*_*i*_(*x*(*t*)) by using *f*_*i*_ = ln *F*_*i*_ and by replacing integrals with respect to continuous-time with sums over discrete-time. The state space for these models is the non-negative cone *C* = [0, ∞)^*n*^ of the *n* dimensional Euclidean space.

### Definitions

A model is <u>dissipative</u> if there exists some density *M >* 0 such that for any initial condition *x*(0) ∈ *C, x*(*t*) enters and remains in the cube [0, *M*]^*n*^ after enough time, i.e., *x*(*t*) ∈ [0, *M*]^*n*^ for *t* sufficiently large. *We assume that the model of interest is dissipative*. Most, but not all, ecological models satisfy this assumption.

Next, we introduce the definition of an attractor for the model. Recall, a compact set *A* ⊂ *C* is <u>invariant</u> if a solution starting in *A* (i.e. *x*(0) ∈ *A*) remains in *A* for all time *t* i.e. *x*(*t*) ∈ *A* for all *t*. The basin of attraction ℬ (*A*) of *A* is the set of initial conditions *x*(0) ∈ *C* such that the solution *x*(*t*) converges to *A* as *t* → ∞, i.e., lim_*t*→∞_ dist(*x*(*t*), *A*) = 0. The word attractor has several meanings depending on the context (Milnor, 1985). For our purposes, a compact set *A* is an <u>attractor</u> if two conditions hold: (i) all points in *A* are “visited” by a single trajectory in forward time (i.e., there exists an initial condition *x*(0) such that 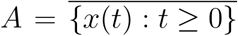) and (ii) the Lebesgue measure of ℬ (*A*) is positive, i.e., there is a positive probability of being in the basin of attraction when randomly choosing an initial condition in [0, *M*]^*n*^ with respect to the uniform distribution. The first condition ensures that the attractor is indecomposable in a certain sense i.e. you can get anywhere on the attractor by following a single trajectory forward in time.

We say *A* is a <u>community attractor</u> if there exists a subset *S* ⊂ {1, 2, …, *n*} of species such that for every *x* ∈ *A, x*_*i*_ *>* 0 if and only if *i* ∈ *S*, i.e., only the species in *S* have positive densities in *A*. We say *S* is the <u>species support</u> of *A*. If *A* is not a community attractor, we say that it is a <u>non-community attractor</u>. By definition, this only occurs if there are different points, say *x, y*, in *A* such that *x*_*i*_ *>* 0 and *y*_*i*_ = 0 for some species *i*. Non-community attractors most naturally arise when there is an attracting heteroclinic cycle, such as in rockpaper-scissors dynamics (May and Leonard, 1975), where community trajectories indefinitely oscillate between different species configurations in an intermittent way.

In order to ensure that invasion growth rates are well defined, we need a key concept from modern dynamical systems: a physical measure also known as Sinai-Bowen-Ruelle (SBR) measure (Young, 1995). A <u>physical measure</u> for an attractor *A* is a probability measure *µ*(*dx*) such that for any continuous function *h* : *C* → ℝ (i.e. “an observable”)

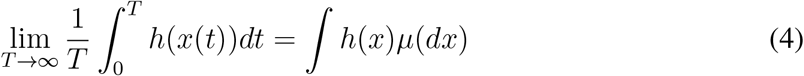

for almost every initial condition *x*(0) ∈ ℬ (*A*) i.e., holds with probability one for a randomly chosen initial condition in ℬ (*A*). In words, equation (4) says “the long-term time average of the observable *h* along the community trajectory equals the spatial average with respect to *µ*(*dx*).” Equivalently, time averages equal ensemble averages. Importantly, for our purposes, the existence of a physical measure implies that there is a well-defined average per-capita growth rate by choosing *h* = *f*_*i*_:

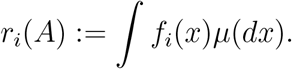

For a community attractor *A*, Schreiber (2000) proved that these average per-capita growth rates are zero for species supported by the attractor, i.e. *r*_*i*_(*A*) = 0 for species *i* that have positive densities in *A*. The average per-capita growth rates corresponding to the non-supported (i.e. “missing”) species are <u>invasion growth rates</u>. *We assume that these invasion growth rates are non-zero*. This assumption has been proven to hold for the “typical” model (Schreiber, 2000; Hening et al., 2022).

#### The Palis Conjecture

With the definitions in hand, we can describe one of the most important conjectures in dynamical systems: the Palis conjecture (Palis, 2000). This conjecture states that the “typical” dynamical system has a finite number of attractors, each of them supporting a physical measure. Formally, “typical” means an open and dense set of models. Practically, this means that unless there is some degenerate dynamical behavior of your model (e.g. the parameters of the model are at a bifurcation value), the conjecture applies to your model. During the past 26 years, there has been significant progress in proving this conjecture for one-dimensional maps, three dimensional ordinary differential equations, and three dimensional stochastic differential equations (Palis, 2000, 2008; Crovisier and Yang, 2015; Crovisier and Pujals, 2015; Hening et al., 2022). The general consensus in the dynamical systems literature is that this conjecture is true.

We present a slight modification of the conjecture for ecological models consistent with the results for stochastic differential equations (Hening et al., 2022). This modification is necessary because the extinction sets (e.g. *x*_1_ = 0 or *x*_2_ = *x*_3_ = 0) are invariant for the dynamics. The standard Palis conjecture does not account for the fact that in ecological models, these extinction boundaries can support robust heteroclinic cycles (e.g. non-community attractors) that persist under perturbation. For example, robust rock-paper-scissor intransitivities can occur along the extinction boundary (May and Leonard, 1975). The modified Palis Conjecture states that the typical ecological dynamical system satisfies two properties:

##### Property 1

**Finite number of attractors** There exists a finite number of attractors *A*_1_, …, *A*_*k*_

for the model restricted to each subset of species (including the full pool of species). The union of the basins of attraction ∪_*i*_ℬ (*A*_*i*_) has the full Lebesgue measure. That is, with probability one, a randomly chosen initial condition in [0, *M*]^*n*^ converges to one of these attractors.

##### Property 2

**Existence of physical measures** Each of the community attractors supports a physical measure.

The only difference from the Palis conjecture is that the non-community attractors need not support a physical measure. Indeed, the lack of physical measures is known for the rock paper scissor model of May and Leonard (1975). When the heteroclinic cycle (i.e., a non-community attractor) is attracting for this model, the time averages of per-capita growth rates (or any nontrivial continuous function) do not converge for any initial conditions in their basin of attraction (Gaunersdorfer and Hofbauer, 1995; Hofbauer and Sigmund, 1998).

#### Numerical implications

The two properties of the modified Palis conjecture have several important implications for the computational identification of community attractors and the calculation of invasion growth rates. Property 1 implies that any randomly chosen initial condition converges (with probability one) to an attractor of the system. Therefore, all attractors can be identified by simulating the model for sufficiently many randomly chosen initial conditions, e.g., what was done for the predator-mediated coexistence example. Property 2 (existence of physical measures) implies that the invasion growth rate can be uniquely defined for community attractors. In addition, for a randomly chosen initial condition in its basin of attraction,

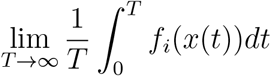

converges (with probability one) to the invasion growth rate given by

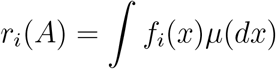

where *µ*(*dx*) is the physical measure associated with the attractor. Thus, simulating the model from random initial conditions will (with probability one) convergence to an attractor, and the time-averaged per-capita growth rates for missing species along the resulting trajectory will (with probability one) converge to the invasion growth rates associated with this attractor. Taken together, simulating the model for sufficiently many random initial conditions not only identifies all the attractors of the system but also allows one to compute the invasion growth rates for all the community attractors, thereby constructing the community disassembly graph. Indeed, these implications provide a mathematical justification for the numerical approach taken by many computationally based approaches to Modern Coexistence Theory (Adler et al., 2010; Ellner et al., 2016; Hallett et al., 2018; Ellner et al., 2019; Shoemaker et al., 2020).

#### The (generalized) community disassembly graph

To define the community disassembly graph, we assume that the (modified) Palis conjecture holds for the model and the model restricted to any subset of species. We also assume, initially, that all attractors are community attractors (see the end of this Appendix for a possible extension). Under these assumptions, the vertices of the community disassembly graph correspond to the finite number of community attractors.

To define the directed edges between these community attractors, let *A* be a community attractor that supports the species in *S*_*A*_ ⊂ {1, 2, …, *k*}. Let *x*(0) ∈ ℬ (*A*) be the community state immediately before the loss of species *i*. After setting *x*_*i*_(0) to zero, *x*(*t*) will converge (with probability one with respect to the choice of *x*(0)) to a community attractor *B* supporting some subset *T* ⊂ *S* \ {*i*} of species that exclude species *i*. The community attractor *B* is a −*I* <u>community for the community attractor</u> *A* and, consequently, there is a directed edge 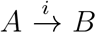 from *A* to *B*. Identifying the −*i* communities for all attractors determines all directed edges of the community disassembly graph.

The following proposition characterizes the −*i* communities using invasion growth rates. The key intuition is that if a species goes extinct following the removal of species *i*, then its long-term average per-capita growth rate along the trajectory to the new attractor must be negative. The proof is based on a related result from (Schreiber, 2025).

##### Proposition

Let *B* be a community attractor that corresponds to a −*i* community for the set of species *S* ⊂ {1, …, *n*} with *i* ∈ *S*. Let *T* ⊂ *S*\{*i*} be the species supported by this attractor, i.e., the species with positive densities in *B*. Then *r*_*j*_(*B*) < 0 for all *j* ∈ *S* \ {*T* ∪ \{*i*}}.

*Proof*. Let *µ*(*dx*) be the physical measure associated with *B* and *x*(0) be an initial condition in the basin of attraction of *B* for which (i) *x*_*j*_(0) *>* 0 for all *j* ∈ *S* \ {*i*} and (ii) 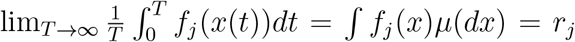 (*B*) for all *j*. Let *j* ∈ *S* \ {*T* ∪ {*i*}}. By the Fundamental Theorem of Calculus and the Chain Rule,

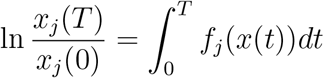

As *j* does not lie in the support of *B* and lim_*T* →∞_ *x*_*j*_(*T*) = 0, we have

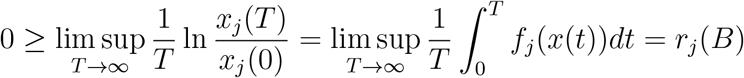

By our assumption that invasion growth rates are non-zero, it follows that *r*_*j*_(*B*) *<* 0. □

To extend the definition of the community disassembly graph to models with non-community attractors, we need to extend our definition of a −*i* community of an attractor *A*. To this end, let *A* be a community attractor that supports the species in *S* ⊂ {1, 2, …, *n*} with *i* ∈ *S*. Let *x*(0) ∈ ℬ (*A*) be the state of the community immediately before the loss of species *i*. After setting *x*_*i*_(0) to zero, *x*(*t*) will converge (with probability one with respect to the choice of *x*(0)) to an attractor *B*. If *B* is a community attractor, it is still a −*i* community as previously defined. However, if *B* is a non-community attractor, then we conjecture that it can be viewed as a heteroclinic cycle involving a finite number of community attractors, say *B*_1_, *B*_2_, …, *B*_*k*_. As *x*(*t*) cycles between these community attractors, the densities of species not supported by any given *B*_*j*_ become arbitrarily low during some parts of the cycle. When demographic stochasticity is present, these low densities make extinction events increasingly likely, ultimately causing the system to converge to one of the community attractors *B*_1_, …, *B*_*k*_ in finite time. For this reason, we propose that (i) each of the community attractors *B*_1_, …, *B*_*k*_ is a −*i* community of the original community attractor *A*, and (ii) all *k* directed edges 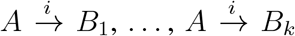 are included in the community disassembly graph. This provides an extension of the community disassembly graph to non-community attractors.

## B Parameters and coexistence of the annual plant model

### B.1 Model and Parameters

The annual plant model from Van Dyke et al. (2022) is

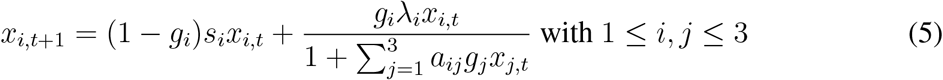

where *x*_*i*_ represents the seed density of species *i*.

Let *A* be the matrix of competition coefficients *a*_*ij*_, 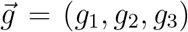 be the vector of germination rates, 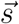 be the vector of seed survival probabilities, and 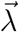 be the vector of maximum birth rates. The parameters from Van Dyke et al. (2022) are

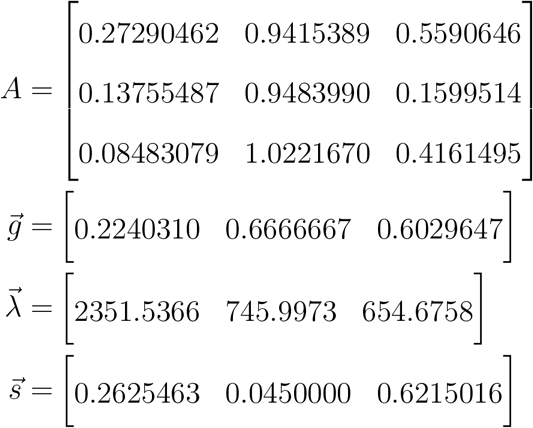

where species 1, 2, and 3 are *A. wrangelianus, H. murinum*, and *P. erecta*, respectively.

### B.2 Solving for attractors of the annual plant model

To identify all the community attractors of the annual plant model (5), we analyze the two subsystems of species using the theory of monotone dynamical systems (Smith, 1995), specifically results for competitive planar maps (Smith, 1998). Then we use invasion graphs (Hofbauer and Schreiber, 2022; Schreiber, 2025) to verify the existence of an attractor that supports all three species.

Ultimately, this analysis only requires solving for the equilibria of (5) and calculating the IGRs at these equilibria. An equilibrium that supports a subset of species *S* ⊂ {1, 2, 3} must satisfy

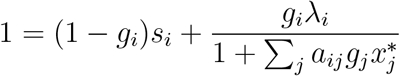

for all species *i* ∈ *S*. Equivalently, it must satisfy the following set of linear equations that are easily solved numerically:

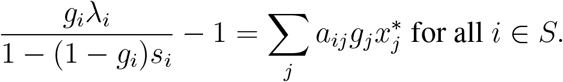

#### B.2.1 Global dynamics of species pairs

We begin by characterizing the dynamics of any pair of species. Without loss of generality, say *i* = 1, 2. Define the update mapping *P*(*x*) = (*P*_1_(*x*), *P*_2_(*x*)) by

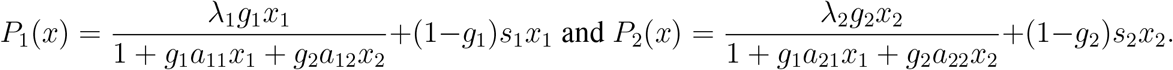

We use results from (Smith, 1998) to characterize the global dynamics. This requires using the theory of monotone maps. The relevant terminology is described in Smith (1995, 1998). First, we verify that (5) with two species (*i* = 1, 2) is strictly competitive in the positive quadrant (0, ∞)^2^. By Proposition 2.1 in (Smith, 1998) it is sufficient to show that *P*_1_ (respectively, *P*_2_) is a strictly increasing function of *x*_1_ and a strictly decreasing function of *x*_2_ (respectively, strictly increasing in *x*_2_ and strictly decreasing in *x*_1_. To this end, we compute the partial derivatives of *F*_1_ and *F*_2_ with respect to *x*_1_ and *x*_2_:

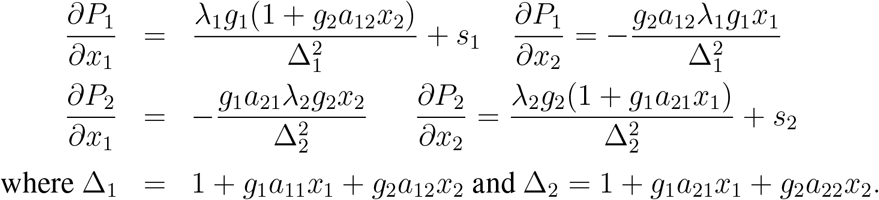

where Δ_1_ = 1 + *g*_1_*a*_11_*x*_1_ + *g*_2_*a*_12_*x*_2_ and Δ_2_ = 1 + *g*_1_*a*_21_*x*_1_ + *g*_2_*a*_22_*x*_2_.

As the partial derivatives 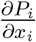 are strictly positive in [0, ∞)^2^ for *i* = 1, 2, the functions *F*_*i*_ increase strictly in *x*_*i*_ in [0, ∞)^2^. As the partial derivatives 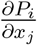 for *i*≠ *j* are negative in (0, ∞)^2^, the functions *F*_*i*_ strictly decrease in *x*_*j*_ (*j* ≠*i*) in the positive quadrant (0, ∞)^2^. Hence, by Proposition 2.3 in (Smith, 1998), (5) is strictly competitive in the positive quadrant.

Next, we verify hypothesis (H+) in (Smith, 1998) for *P* on *A* = (0, ∞)^2^. This hypothesis consists of four conditions (a)–(d). Condition (a) holds immediately for the competitive ordering <_*K*_ on *A*. Condition (b) requires that the determinant of the Jacobian matrix

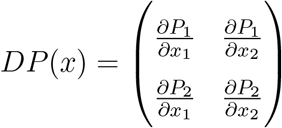

is positive in (0, ∞)^2^. The following calculation ensures that this condition is satisfied:

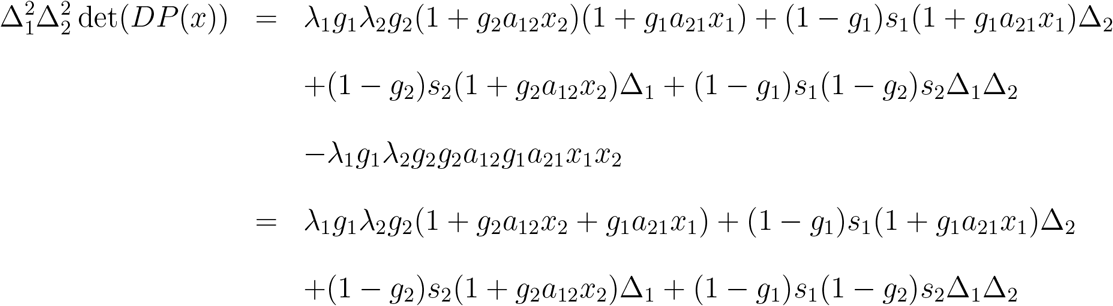

The third condition (c) requires that the diagonal entries of *DP*(*x*) be strictly positive and the off-diagonal elements be strictly negative on (0, ∞)^2^. This follows immediately from our partial derivative calculations. The final condition (d) requires that *P* is injective in (0, ∞)^2^. As det(*DP*(*x*)) > 0 on [0, ∞)^2^ and [0, ∞)^2^ is connected, all points in (0, ∞)^2^ have the same number of preimages. As the only preimage of (0, 0) is (0, 0), it follows that *P* is injective.

Having verified these conditions, Corollary 4.4 in (Smith, 1998) implies that all trajectories of (B) for *i* = 1, 2 converge to an equilibrium. To understand which initial conditions converge to which equilibria, we need to solve the equilibria and check the invasion criteria at these equilibria. As we describe below, there is for our parameters at most one equilibrium for any subset of species. Moreover, for any pair of species in the parameterized model, only one of the two mutual invasibility conditions holds. Either both species have a positive invasion growth rate at the other species equilibrium or the invasion growth rates are of opposite sign. When the former occurs, the single species equilibria are unstable (saddles), and there is a unique equilibrium, call it *x*^∗^, supporting the other two species, i.e. 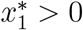 and 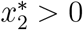. Therefore, any trajectory where initially both species have positive densities converges to *x*^∗^ i.e. *x*^∗^ is globally stable. Alternatively, suppose that the invasion growth rates have opposite signs. Without loss of generality, say the invasion growth rate of species 2 is negative at the equilibrium (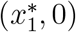, 0) of species 1. Then there is no equilibrium supporting both species and the extinction equilibrium (0, 0) and the species 2 equilibrium (0, 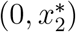) are unstable, while the species 1 equilibrium is stable. Therefore, Corollary 4.4 implies that all trajectories at which species 1 is initial present converge to (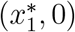, 0). These results fully characterize the dynamics of all the two species pairs in the three species parameterized model.

#### B.2.2 Three species coexistence

We use the invasion graph approach (Figure 7) to verify coexistence. This approach was introduced for differential equations in Hofbauer and Schreiber (2022). Calculating the equilibria as detailed below for all single and two species subsystems, as described below, and using the convergence results for the single and two species subsystems, one can verify that the invasion graph, as shown in Figure 7, is acyclic and that the invasion growth rate of species *i* for *i* = 1, 2, 3 is positive at the −i equilibria. Hence, Theorem 4.8 in Schreiber (2025) implies that all three species coexist in the sense of permanence and, therefore, at a feasible attractor. Numerical simulations suggest that permanence occurs via a global stable equilibrium.

**Figure 7:**
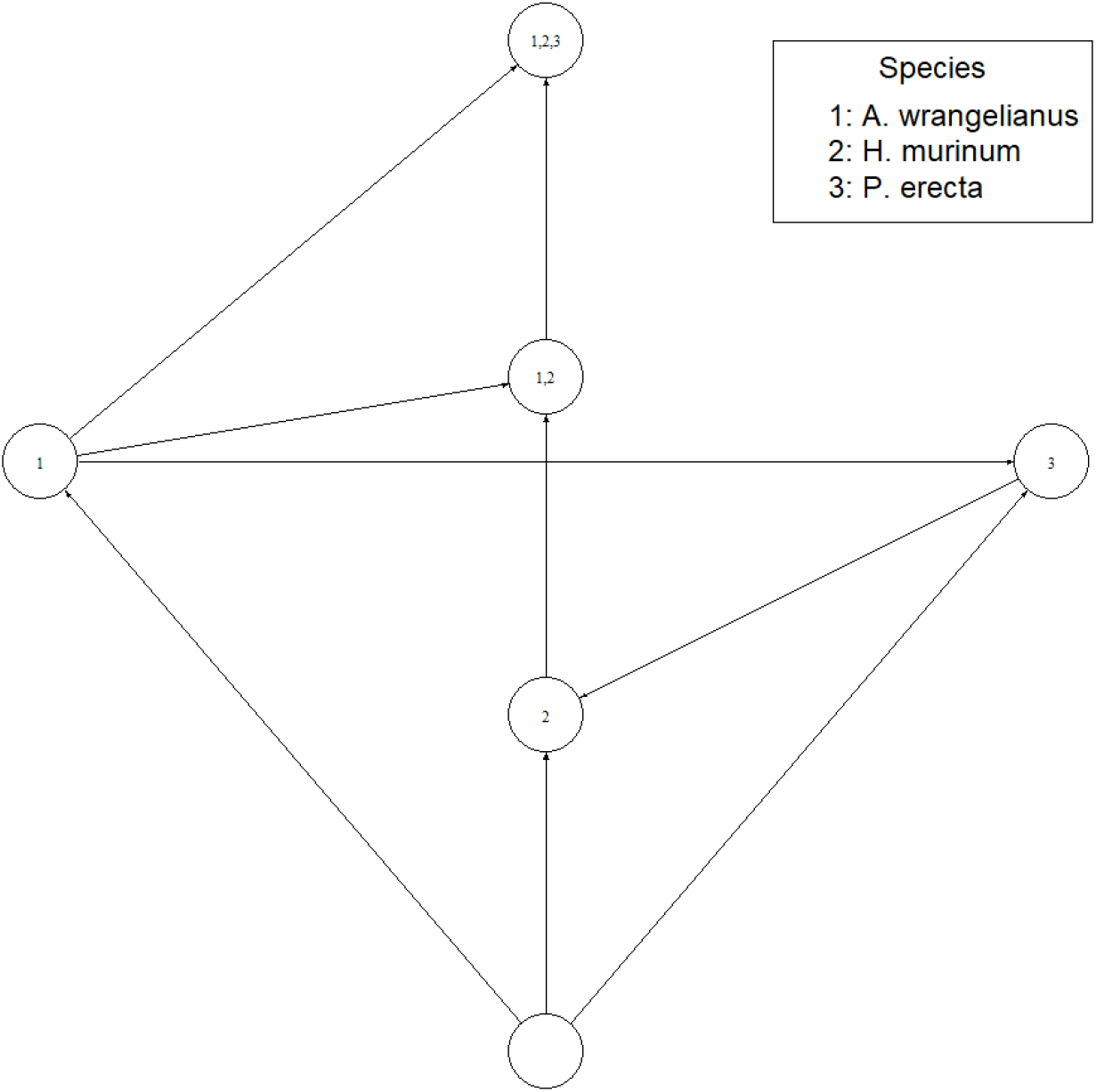
Invasion graph of the annual plant model from Van Dyke et al. (2022). Nodes represent feasible equilibria while directed edges represent changes in community composition as a result of species invasion.

**Figure 8:**
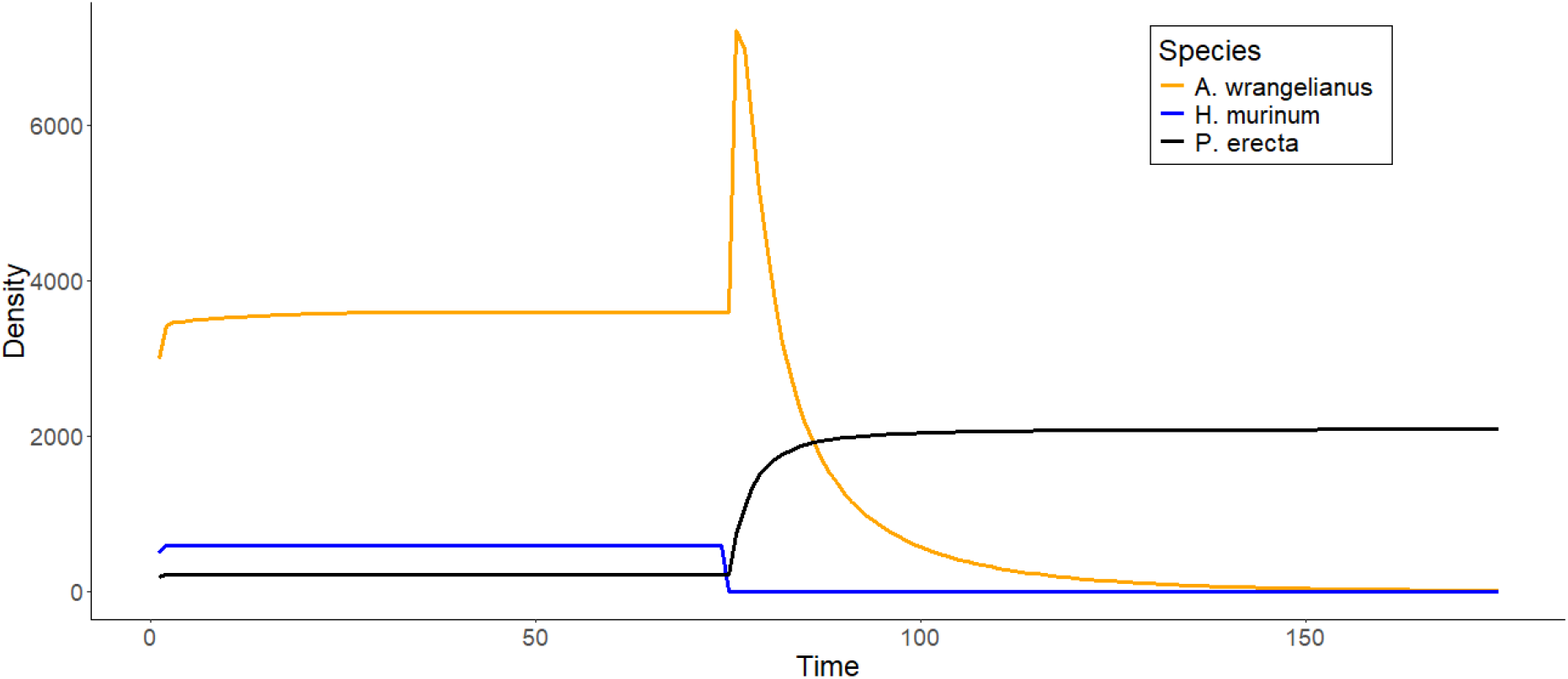
Simulation of community dynamics following extinction of *H. murinum* at time step 75. Note that this leads to a secondary extinction of *A. wrangelianus*, matching the -*H. murinum* predicted in the community disassembly graph in Figure 1A.

## C Alternative secondary extinction in annual plant model

Revisiting the annual plant model, there exists another secondary extinction event in the full community. As shown in Figure 1A & 8, extinction of *H. murinum* leads to the secondary extinction of *A. wrangelianus*. As the procedure follows, we will (1) understand why *A. wrangelianus* is excluded by *P. erecta* and (2) why it can invade the community with *P. erecta* and *H. murinum* is present.

Figure 1A shows that the -*A. wrangelianus* community has a secondary extinction event of *P. erecta*. Thus, introduction of *A. wrangelianus* alone cannot restore the pre-disassembled community. Thus, we identify a path from the -*A. wrangelianus community* back to the full community. For this empirical example, there exists only one path from -*A. wrangelianus* community to the full community: invasion of *A. wrangelianus* in -*A. wrangelianus* community that changes the composition to coexistence of *A. wrangelianus* and *H. murinum*, followed by invasion of *P. erecta* to reassemble the system back to the full community (Figure 9).

**Figure 9:**
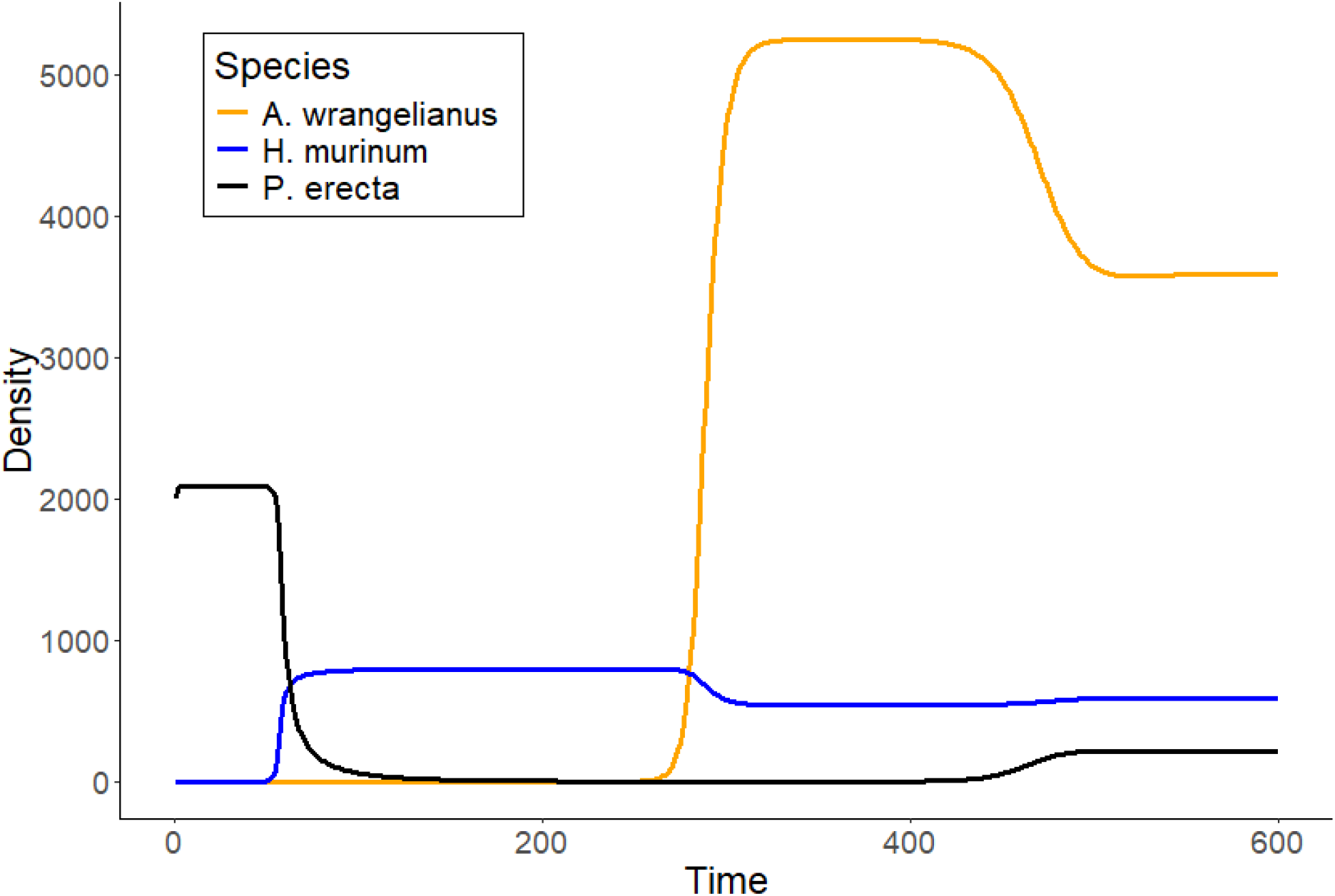
Simulation of dynamics associated with invasion. Invasion of *H. murinum* at time step 50; invasion of *A. wrangelianus* at time step 250; invasion of *P. erecta* at time step 400.

Along this assembly path, *A. wrangelianus* only needs to successfully invade once at the -*A. wrangelianus community* since it is not displaced by subsequent invasions on the assembly path. Thus, we simply decompose the invasion growth rate of *A. wrangelianus* at the -*A. wrangelianus* community to understand why it successfully invades in this community.

As shown in Figure 10, *A. wrangelianus* can invade the −*A. wrangelianus* community, only containing *H. murinum*. Furthermore, since *H. murinum* has a positive invasion growth rate in the community only containing *A. wrangelianus*, we know they coexist through mutual invasibility.

**Figure 10:**
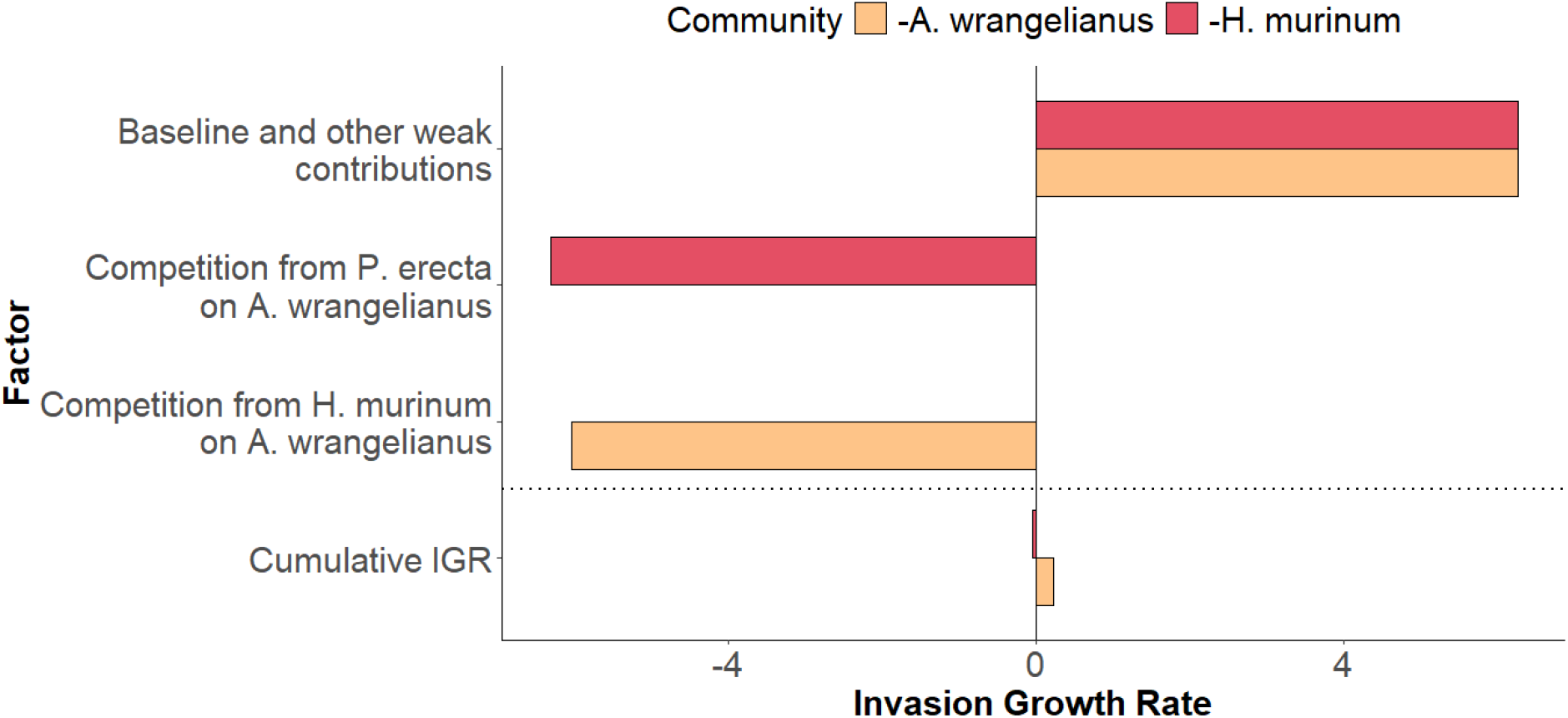
Decomposition of *A. wrangelianus*’ IGR at the −*H. murinum* and −*A. wrangelianus* communities. The sum of all bars of a respective color above the dotted line add to the value of the bar below the dotted line of the respective color. Factors not present in the −*i* community have no contribution to the IGR and thus have no corresponding bar.

**Figure 11:**
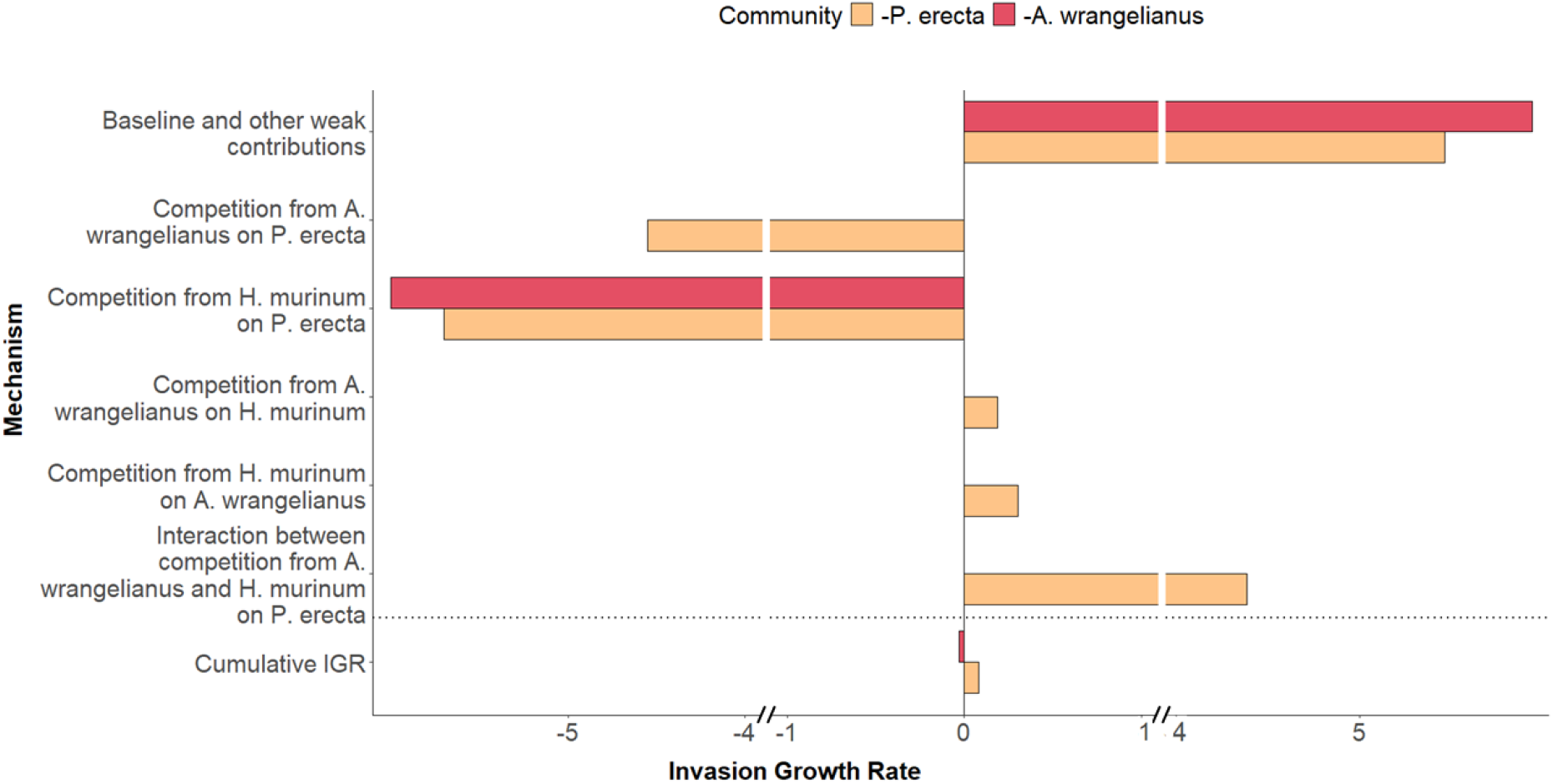
Decomposition of *P. erecta*’s IGR at the −*P. erecta* and −*A. wrangelianus* communities. The sum of all bars of a respective color above the dotted line add to the value of the bar below the dotted line of the respective color. Factors whose value is less than 0.01 were lumped into the baseline bar. Mechanisms not present in the −*i* community have no contribution to the IGR and thus have no bar represented. Note that this decomposition has interactive terms due to the nonlinearity of competition coefficients in the invasion growth rates.

In the disassembled community, *A. wrangelianus* cannot coexist with *P. erecta* since *P. erecta* is a superior competitor to *A. wrangelianus*. However, despite this, we find that *A. wrangelianus* and *H. murinum* can coexist. When *P. erecta* then invades the community with *A. wrangelianus* and *H. murinum*, it invades without displacing *A. wrangelanius* due to *H. murinum*’s competitive pressure on *P. erecta*, leading to coexistence of the full community. Thus, we see that persistence of *A. wrangelianus* in the full community is contingent upon the presence of *H. murinum*.

## D Alternate decomposition of *P. erecta*’s IGR

## E Parameters and coexistence for grassland community model

### E.1 Model and Parameters

The model for the grassland community in Geijzendorffer et al. (2011) discussed in the “Facilitation” section of the main text is

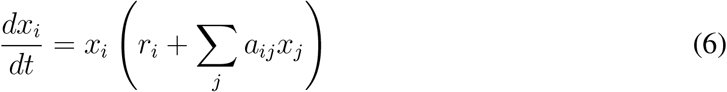

where *x*_*i*_ is the biomass of species *i, r*_*i*_ is the intrinsic growth rate of species *i*, and *a*_*ij*_ is the per-capita competitive effect of species *j* on *i*.

Let *A* be the matrix of competition coefficients *a*_*ij*_ and 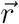 be the vector of intrinsic growth rates *r*_*i*_. From Geijzendorffer et al. (2011), the model parameters for *A. stolonifera, L. perenne, P. pratense, T. pratense*, and *T. repens* are respectively

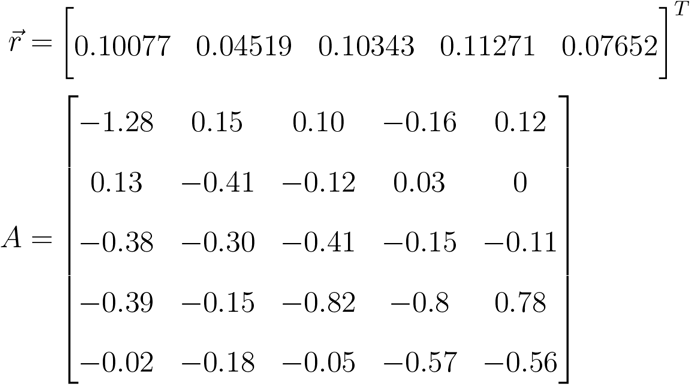

### E.2 Solving for attractors of the grassland community model

To determine the coexistence states of the system, we computed the invasion graph (Hofbauer and Schreiber, 2022) which is shown in Figure 12. This invasion graph is acyclic and all nodes satisfy the permanence condition. Hence, all nodes correspond to not only local but global attractors.

**Figure 12:**
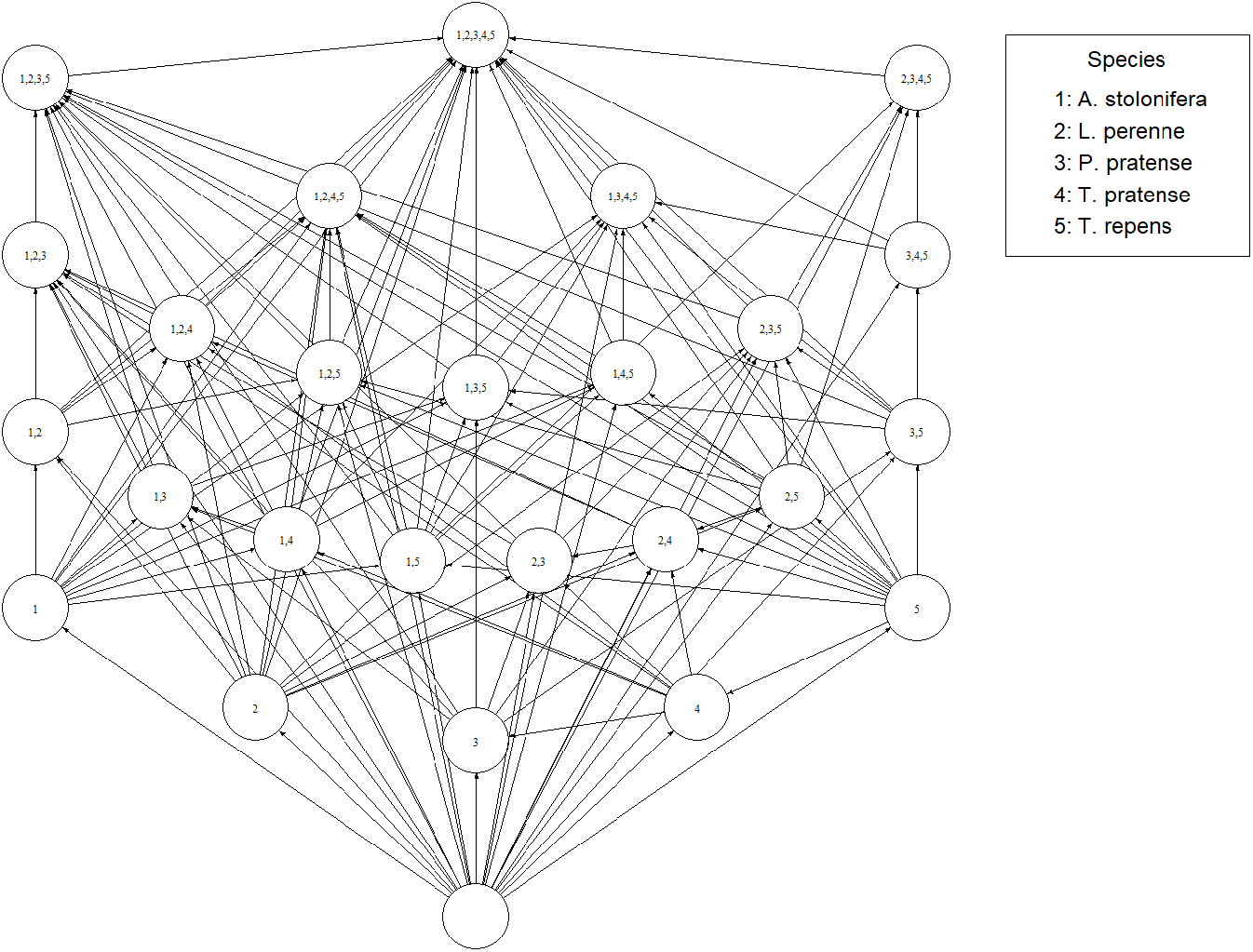
Invasion graph of the grassland community from Geijzendorffer et al. (2011). Nodes represent feasible equilibria while directed edges represent changes in community composition as a result of species invasion.

## F Parameters for diamond model

Here, we use the same parametrization as Vasseur and Fox (2007) and Shoemaker et al. (2020) for the model of keystone predation described in the “Predation” section of the main text.

| Parameter | Description | Value |
| --- | --- | --- |
| $r$ | Resource intrinsic rate of growth | 1.0 |
| $K$ | Resource carrying capacity | 1.0 |
| $J_{C_1}$ | Competitor 1's ingestion rate | 0.8036 |
| $J_{C_2}$ | Competitor 2's ingestion rate | 0.7 |
| $J_P$ | Predator ingestion rate | 0.4 |
| $M_{C_1}$ | Competitor 1's intrinsic mortality rate | 0.4 |
| $M_{C_2}$ | Competitor 2's intrinsic mortality rate | 0.2 |
| $M_P$ | Predator's intrinsic mortality rate | 0.08 |
| $R_{0_1}$ | Competitor 1's half saturation constant of the resource | 0.16129 |
| $R_{0_2}$ | Competitor 2's half saturation constant of the resource | 0.9 |
| $C_0$ | Predator's half saturation constant of the competitors | 0.5 |
| $\Omega_{PC_1}$ | Predator's prey preference for competitor 1 | 0.92 |
| $\Omega_{C_1R}$ | Competitor 1's preference coefficient for the resource | 1.0 |
| $\Omega_{C_2R}$ | Competitor 2's preference coefficient for the resource | 0.98 |

## Notes

https://github.com/biologyjoe87/MCT_disassembly

### Competing Interest Statement

The authors have declared no competing interest.

### Summary of Updates

We have updated the methods and discussion of the manuscript. Additionally, we have added more mathematical details into the supplement.

## References

K. C. Abbott, C. M. Heggerud, Y.-C. Lai, A. Morozov, S. Petrovskii, K. Cuddington, and A. Hastings. When and why ecological systems respond to the rate rather than the magnitude of environmental changes. Biological Conservation, 292:110494, 2024.

P. B. Adler, S. P. Ellner, and J. M. Levine. Coexistence of perennial plants: an embarrassment of niches. Ecology letters, 13(8): 1019–1029, 2010.

R. A. Armstrong and R. McGehee. Competitive exclusion. The American Naturalist, 115(2): 151–170, 1980.

C. E. Aslan, E. S. Zavaleta, B. Tershy, and D. Croll. Mutualism disruption threatens global plant biodiversity: a systematic review. PloS one, 8(6):e66993, 2013.

M. S. Bane, M. J. Pocock, and R. James. Effects of model choice, network structure, and interaction strengths on knockout extinction models of ecological robustness. Ecology and evolution, 8(22): 10794–10804, 2018.

G. Barabás, R. D’Andrea, and S. M. Stump. Chesson’s coexistence theory. Ecological monographs, 88(3): 277–303, 2018.

I. Bartomeus, S. Saavedra, R. P. Rohr, and O. Godoy. Experimental evidence of the importance of multitrophic structure for species persistence. Proceedings of the National Academy of Sciences, 118(12):e2023872118, 2021.

J. Bascompte and D. B. Stouffer. The assembly and disassembly of ecological networks. Philosophical Transactions of the Royal Society B: Biological Sciences, 364(1524): 1781–1787, 2009.

J. F. Brodie, C. E. Aslan, H. S. Rogers, K. H. Redford, J. L. Maron, J. L. Bronstein, and C. R. Groves. Secondary extinctions of biodiversity. Trends in ecology & evolution, 29(12): 664–672, 2014.

P. Chesson. Multispecies competition in variable environments. Theoretical population biology, 45(3): 227–276, 1994.

A. T. Clark, L. G. Shoemaker, J.-F. Arnoldi, G. Barabás, R. Germain, O. Godoy, L. Hallett, C. Karakoç, S. Saavedra, and S. Schreiber. A practical guide to quantifying ecological coexistence. 2024.

R. K. Colwell, R. R. Dunn, and N. C. Harris. Coextinction and persistence of dependent species in a changing world. Annual Review of Ecology, Evolution, and Systematics, 43(1): 183–203, 2012.

S. Crovisier and E. R. Pujals. Essential hyperbolicity and homoclinic bifurcations: a dichotomy phenomenon/mechanism for diffeomorphisms. Inventiones mathematicae, 201(2): 385–517, 2015.

S. Crovisier and D. Yang. On the density of singular hyperbolic three-dimensional vector fields: a conjecture of palis. Comptes Rendus. Mathématique, 353(1): 85–88, 2015.

A. Curtsdotter, A. Binzer, U. Brose, F. de Castro, B. Ebenman, A. Eklöf, J. O. Riede, A. Thierry, and B. C. Rall. Robustness to secondary extinctions: comparing trait-based sequential deletions in static and dynamic food webs. Basic and Applied Ecology, 12(7): 571–580, 2011.

J. A. Dunne, R. J. Williams, and N. D. Martinez. Food-web structure and network theory: the role of connectance and size. Proceedings of the National Academy of Sciences, 99(20): 12917–12922, 2002a.

J. A. Dunne, R. J. Williams, and N. D. Martinez. Network structure and biodiversity loss in food webs: robustness increases with connectance. Ecology letters, 5(4):558–567, 2002b.

J. A. Dunne, R. J. Williams, and N. D. Martinez. Network structure and robustness of marine food webs. Marine Ecology Progress Series, 273: 291–302, 2004.

B. Ebenman and T. Jonsson. Using community viability analysis to identify fragile systems and keystone species. Trends in Ecology & Evolution, 20(10): 568–575, 2005.

B. Ebenman, R. Law, and C. Borrvall. Community viability analysis: the response of ecological communities to species loss. Ecology, 85(9): 2591–2600, 2004.

K. F. Edwards and S. J. Schreiber. Preemption of space can lead to intransitive coexistence of competitors. Oikos, 119(7): 1201–1209, 2010.

A. J. Eichenwald and J. M. Reed. An expanded framework for community viability analysis. Bioscience, 71(6): 626–636, 2021.

A. Eklöf and B. Ebenman. Species loss and secondary extinctions in simple and complex model communities. Journal of animal ecology, pages 239–246, 2006.

S. P. Ellner, R. E. Snyder, and P. B. Adler. How to quantify the temporal storage effect using simulations instead of math. Ecology letters, 19(11): 1333–1342, 2016.

S. P. Ellner, R. E. Snyder, P. B. Adler, and G. Hooker. An expanded modern coexistence theory for empirical applications. Ecology letters, 22(1): 3–18, 2019.

S. P. Ellner, R. E. Snyder, P. B. Adler, and G. Hooker. Toward a “modern coexistence theory”‘ for the discrete and spatial. Ecological Monographs, 92(4):e1548, 2022.

C. Emary and D. Evans. Can a complex ecosystem survive the loss of a large fraction of its species? a random matrix theory of secondary extinction. Oikos, 130(9): 1512–1522, 2021.

A. Gaunersdorfer and J. Hofbauer. Fictitious play, shapley polygons, and the replicator equation. Games and Economic Behavior, 11(2): 279–303, 1995.

G. F. Gause. The struggle for existence. 1934.

I. Geijzendorffer, W. Van der Werf, F. Bianchi, and R. Schulte. Sustained dynamic transience in a lotka–volterra competition model system for grassland species. Ecological Modelling, 222(15): 2817–2824, 2011.

R. M. Germain, J. L. Williams, D. Schluter, and A. L. Angert. Moving character displacement beyond characters using contemporary coexistence theory. Trends in Ecology & Evolution, 33(2): 74–84, 2018.

P. W. Glynn. Some physical and biological determinants of coral community structure in the eastern pacific. Ecological monographs, 46(4): 431–456, 1976.

N. A. Graham, P. Chabanet, R. D. Evans, S. Jennings, Y. Letourneur, M. Aaron MacNeil, T. R. McClanahan, M. C. Öhman, N. V. Polunin, and S. K. Wilson. Extinction vulnerability of coral reef fishes. Ecology letters, 14(4): 341–348, 2011.

T. N. Grainger, A. D. Letten, B. Gilbert, and T. Fukami. Applying modern coexistence theory to priority effects. proceedings of the national academy of sciences, 116(13):6205–6210, 2019a.

T. N. Grainger, J. M. Levine, and B. Gilbert. The invasion criterion: a common currency for ecological research. Trends in ecology & evolution, 34(10):925–935, 2019b.

L. M. Hallett, E. C. Farrer, K. N. Suding, H. A. Mooney, and R. J. Hobbs. Tradeoffs in demographic mechanisms underlie differences in species abundance and stability. Nature Communications, 9(1): 5047, 2018.

A. Hastings, K. C. Abbott, K. Cuddington, T. Francis, G. Gellner, Y.-C. Lai, A. Morozov, S. Petrovskii, K. Scranton, and M. L. Zeeman. Transient phenomena in ecology. Science, 361(6406):eaat6412, 2018.

A. Hening, D. H. Nguyen, and S. J. Schreiber. A classification of the dynamics of three-dimensional stochastic ecological systems. The Annals of Applied Probability, 32(2): 893–931, 2022.

J. Hofbauer. Heteroclinic cycles in ecological differential equations. Equadiff 8, pages 105–116, 1994.

J. Hofbauer and S. J. Schreiber. Permanence via invasion graphs: incorporating community assembly into modern coexistence theory. Journal of mathematical biology, 85(5): 54, 2022.

J. Hofbauer and K. Sigmund. Evolutionary games and population dynamics. Cambridge university press, 1998.

J. Jackson and L. Buss. Alleopathy and spatial competition among coral reef invertebrates. Proceedings of the National Academy of Sciences, 72(12): 5160–5163, 1975.

F. Jordán. Keystone species and food webs. Philosophical Transactions of the Royal Society B: Biological Sciences, 364(1524): 1733–1741, 2009.

P.-J. Ke and A. D. Letten. Coexistence theory and the frequency-dependence of priority effects. Nature Ecology & Evolution, 2(11): 1691–1695, 2018.

R. Kehoe, E. Frago, and D. Sanders. Cascading extinctions as a hidden driver of insect decline. Ecological Entomology, 46(4): 743–756, 2021.

B. Kerr, M. A. Riley, M. W. Feldman, and B. J. Bohannan. Local dispersal promotes biodiversity in a real-life game of rock–paper–scissors. Nature, 418(6894): 171–174, 2002.

R. E. Langendorf, J. A. Estes, J. C. Watson, M. C. Kenner, B. B. Hatfield, M. T. Tinker, E. Waddle, M. L. DeMarche, and D. F. Doak. Dynamic and context-dependent keystone species effects in kelp forests. Proceedings of the National Academy of Sciences, 122(10): e2413360122, 2025.

M. Letnic, F. Koch, C. Gordon, M. S. Crowther, and C. R. Dickman. Keystone effects of an alien top-predator stem extinctions of native mammals. Proceedings of the Royal Society B: Biological Sciences, 276(1671): 3249–3256, 2009.

J. M. Levine, J. Bascompte, P. B. Adler, and S. Allesina. Beyond pairwise mechanisms of species coexistence in complex communities. Nature, 546(7656): 56–64, 2017.

J. Lubchenco. Plant species diversity in a marine intertidal community: importance of herbivore food preference and algal competitive abilities. The American Naturalist, 112(983): 23–39, 1978.

R. MacArthur and R. Levins. The limiting similarity, convergence, and divergence of coexisting species. The american naturalist, 101(921): 377–385, 1967.

M. Manhart and E. I. Shakhnovich. Growth tradeoffs produce complex microbial communities on a single limiting resource. Nature communications, 9(1): 3214, 2018.

R. M. May and W. J. Leonard. Nonlinear aspects of competition between three species. SIAM journal on applied mathematics, 29(2): 243–253, 1975.

K. McCann, A. Hastings, and G. R. Huxel. Weak trophic interactions and the balance of nature. Nature, 395(6704): 794–798, 1998.

J. Memmott, N. M. Waser, and M. V. Price. Tolerance of pollination networks to species extinctions. Proceedings of the Royal Society of London. Series B: Biological Sciences, 271 (1557):2605–2611, 2004.

J. Milnor. On the concept of attractor. Communications in Mathematical Physics, 99(2): 177–195, 1985.

M. L. Moir, P. A. Vesk, K. E. Brennan, D. A. Keith, L. Hughes, and M. A. McCARTHY. Current constraints and future directions in estimating coextinction. Conservation Biology, 24(3): 682–690, 2010.

R. S. Ostfeld and K. LoGiudice. Community disassembly, biodiversity loss, and the erosion of an ecosystem service. Ecology, 84(6): 1421–1427, 2003.

R. T. Paine. Food web complexity and species diversity. The American Naturalist, 100(910): 65–75, 1966.

R. T. Paine. Food webs: linkage, interaction strength and community infrastructure. Journal of animal ecology, 49(3): 667–685, 1980.

R. T. Paine, J. Terborgh, and J. Estes. Food chain dynamics and trophic cascades in intertidal habitats. Trophic cascades: predators, prey, and the changing dynamics of nature, pages 21–35, 2010.

J. Palis. A global view of dynamics and a conjecture on the denseness of finitude of attractors. Astérisque, 261(xiiixiv):335–347, 2000.

J. Palis. Open questions leading to a global perspective in dynamics. Nonlinearity, 21(4):T37, 2008.

A. I. Pastore, G. Barabás, M. D. Bimler, M. M. Mayfield, and T. E. Miller. The evolution of niche overlap and competitive differences. Nature Ecology & Evolution, 5(3): 330–337, 2021.

S. C. Pennings and R. M. Callaway. Impact of a parasitic plant on the structure and dynamics of salt marsh vegetation. Ecology, 77(5): 1410–1419, 1996.

R. Ranjan, T. Koffel, and C. A. Klausmeier. The three-species problem: Incorporating competitive asymmetry and intransitivity in modern coexistence theory. Ecology Letters, 27(4): e14426, 2024.

A. G. Rossberg, A. L. Caskenette, and L.-F. Bersier. Structural instability of food webs and food-web models and their implications for management. Adaptive food webs: stability and transitions of real and model ecosystems, pages 372–383, 2017.

S. Saavedra, R. P. Rohr, J. Bascompte, O. Godoy, N. J. Kraft, and J. M. Levine. A structural approach for understanding multispecies coexistence. Ecological Monographs, 87(3): 470–486, 2017.

S. J. Schreiber. Criteria for cr robust permanence. Journal of Differential Equations, 162(2): 400–426, 2000.

S. J. Schreiber. Coexistence and extinction in flow-kick systems: An invasion growth rate approach. arXiv preprint arXiv:2507.02157, 2025.

S. J. Schreiber and S. Rittenhouse. From simple rules to cycling in community assembly. Oikos, 105(2): 349–358, 2004.

S. J. Schreiber, M. Yamamichi, and S. Y. Strauss. When rarity has costs: coexistence under positive frequency-dependence and environmental stochasticity. Ecology, 100(7):e02664, 2019.

L. G. Shoemaker, A. K. Barner, L. S. Bittleston, and A. I. Teufel. Quantifying the relative importance of variation in predation and the environment for species coexistence. Ecology Letters, 23(6): 939–950, 2020.

H. L. Smith. Monotone dynamical systems: An introduction to the theory of competitive and cooperative systems. Number 41. American Mathematical Society, 1995.

H. L. Smith. Planar competitive and cooperative difference equations. Journal of Difference Equations and Applications, 3(5–6): 335–357, 1998.

K. Soetaert, T. Petzoldt, and R. W. Setzer. Solving differential equations in r: package desolve. Journal of statistical software, 33: 1–25, 2010.

C. Song and J. W. Spaak. Trophic tug-of-war: Coexistence mechanisms within and across trophic levels. Ecology Letters, 27(4):e14409, 2024.

J. W. Spaak and S. J. Schreiber. Building modern coexistence theory from the ground up: the role of community assembly. Ecology Letters, 26(11): 1840–1861, 2023.

J. J. Stachowicz. Mutualism, facilitation, and the structure of ecological communities: positive interactions play a critical, but underappreciated, role in ecological communities by reducing physical or biotic stresses in existing habitats and by creating new habitats on which many species depend. Bioscience, 51(3): 235–246, 2001.

J. J. Stachowicz and M. E. Hay. Mutualism and coral persistence: the role of herbivore resistance to algal chemical defense. Ecology, 80(6): 2085–2101, 1999.

G. Strona and C. J. Bradshaw. Co-extinctions annihilate planetary life during extreme environmental change. Scientific reports, 8(1): 16724, 2018.

Y. Tang, F. Wang, and W. Zhou. Network structure indicators predict ecological robustness in food webs. Ecological Research, 39(5): 766–774, 2024.

D. Tilman. Resource competition and community structure. Number 17. Princeton university press, 1982.

A. Torres, S. E. Kuebbing, K. L. Stuble, S. A. Catella, M. A. Núñez, and M. A. Rodriguez-Cabal. Inverse priority effects: A role for historical contingency during species losses. Ecology Letters, 27(1):e14360, 2024a.

A. Torres, T. Morán-López, M. A. Rodriguez-Cabal, and M. A. Núñez. Inverse priority effects: The order and timing of removal of invasive species influence community reassembly. Journal of Applied Ecology, 61(1):51–62, 2024b.

M. N. Van Dyke, J. M. Levine, and N. J. Kraft. Small rainfall changes drive substantial changes in plant coexistence. Nature, 611(7936): 507–511, 2022.

A. J. Vanbergen, B. A. Woodcock, M. S. Heard, and D. S. Chapman. Network size, structure and mutualism dependence affect the propensity for plant–pollinator extinction cascades. Functional ecology, 31(6): 1285–1293, 2017.

D. A. Vasseur and J. W. Fox. Environmental fluctuations can stabilize food web dynamics by increasing synchrony. Ecology letters, 10(11): 1066–1074, 2007.

A. D. Wallach, W. J. Ripple, and S. P. Carroll. Novel trophic cascades: apex predators enable coexistence. Trends in Ecology & Evolution, 30(3): 146–153, 2015.

J. P. Wright, C. G. Jones, and A. S. Flecker. An ecosystem engineer, the beaver, increases species richness at the landscape scale. Oecologia, 132: 96–101, 2002.

J. D. Yeakel, M. M. Pires, M. A. de Aguiar, J. L. O’Donnell, P. R. Guimarães Jr, D. Gravel, and T. Gross. Diverse interactions and ecosystem engineering can stabilize community assembly. Nature communications, 11(1): 3307, 2020.

L.-S. Young. Ergodic theory of differentiable dynamical systems. NATO ASI Series C Mathematical and Physical Sciences-Advanced Study Institute, 464: 293–336, 1995.

E. Zavaleta, J. Pasari, J. Moore, D. Hernandez, K. B. Suttle, and C. C. Wilmers. Ecosystem responses to community disassembly. Annals of the New York Academy of Sciences, 1162 (1):311–333, 2009.

